# Genomic epidemiological models describe pathogen evolution across fitness valleys

**DOI:** 10.1101/2021.12.16.473045

**Authors:** Pablo Cárdenas, Vladimir Corredor, Mauricio Santos-Vega

## Abstract

Genomics is fundamentally changing epidemiological research. However, systematically exploring hypotheses in pathogen evolution requires new modeling tools. Models intertwining pathogen epidemiology and genomic evolution can help understand processes such as the emergence of novel pathogen genotypes with higher transmissibility or resistance to treatment. In this work, we present Opqua, a flexible simulation framework that explicitly links epidemiology to sequence evolution and selection. We use Opqua to study determinants of evolution across fitness valleys. We confirm that competition can limit evolution in high transmission environments and find that low transmission, host mobility, and complex pathogen life cycles facilitate reaching new adaptive peaks through population bottlenecks and decoupling of selective pressures. The results show the potential of genomic epidemiological modeling as a tool in infectious disease research.

## Introduction

Genomic epidemiology has become a powerful tool in the study and control of infectious disease spread. By sequencing the genomes of pathogens in the field, researchers can access a living ledger of transmission and evolution in real-time, which aids understanding of disease spread and can inform public health policy. Sequencing of pathogen genomes has been used to monitor evolution, trace local chains of transmission, and study the origins of outbreaks in real-time for Ebola (*1–3*), malaria (*4–6*), influenza (*7, 8*), and COVID-19 (*9–19*), among others. An arsenal of tools and initiatives have been developed to harness the full potential of data generated through genomic surveillance (*4, 20–23*).

Despite all its advantages, genomic epidemiology is hampered by its ability to experimentally explore hypotheses beyond observational data. This is a problem shared by most epidemiological research. Fortunately, mathematical and computational modeling provide frameworks for investigating the effect of individual variables through the simulation of null models and perfectly-controlled, targeted interventions, which would otherwise be impossible in practice (*12, 24–26*). However, most traditional modeling frameworks are not well-suited to work with genomic data and the kinds of evolutionary questions unlocked by genomic epidemiology. For example, different computational tools have been developed to simulate the evolution of pathogen genetic sequences (*27–29*), but in these cases, the tools are not coupled to epidemiological models that follow disease spread.

More recently, there has been interest in developing tools to combine epidemiological models with genetic sequence simulators [such as mrc-ide.github.io/SIMPLEGEN/index.html, (*30–33*)]. While these approaches can be powerful tools to answer questions about population genetics, they do not consider the effect of genetic sequences on disease dynamics. This makes them unable to account for selection in pathogen evolution. More general tools are available to simulate the evolution of organism populations in forward-time (*34–36*). Although these may be adapted to use in epidemiology, this requires significant work and setup, especially when tailoring to different kinds of disease transmission and intervention. Further recent work has been done to explore evolving epidemiological models in a custom, disease-specific context (*37*). Nevertheless, there currently is no out-of-the-box solution for building flexible, easy-to-use simulations of disease spread with pathogens capable of evolving and influencing their epidemiology through natural selection.

Simulating selection and evolution becomes particularly pressing as we ask questions about how epidemiology affects the appearance of advantageous traits within a pathogen population. Specifically, we are interested in how epidemiological contexts can shape competition and clonal interference between pathogens to hamper the evolution of novel traits separated by fitness valleys. Significant work has examined the evolutionary dynamics of crossing fitness valleys, a process termed stochastic tunneling (*38–41*). Other work has examined the environmental factors that shape these adaptive landscapes (*42*). Still, quantitative research into stochastic tunneling in the context of a spreading infection has only recently become an object of study (*37*). This is notable given the interest in the evolutionary biology of infectious diseases sparked by the COVID-19 pandemic. The evolutionary trajectory of its causing agent, SARS-CoV-2, has brought interest into the role of epidemiology in shaping evolutionary trajectories across fitness valleys. Thus, epidemiology becomes crucial in understanding future viral evolution and vaccine effectiveness. The emergence of novel, evolutionarily distant SARS-CoV-2 variants such as Omicron is a concerning testament to the importance of these questions, as this new variant bears multiple compensatory mutations and epistatic interactions that individually decrease fitness, but have a synergistic effect on transmission (*43, 44*). Studying the eco-epidemiological determinants of evolution is also key to understanding the emergence of drug resistance in diseases such as tuberculosis and malaria, where fitness costs of resistant mutations are offset by a succession of compensatory mutations (*45, 46*).

In light of this, we developed Opqua (github.com/pablocarderam/opqua) as an epidemiological modeling framework for pathogen population genetics and evolution. We apply Opqua to explore how different aspects of the epidemiological environment and infection biology constrain or enhance the ability of pathogens to escape local fitness peaks and explore new evolutionary space in adaptive landscapes.

### A computational modeling framework for genomic epidemiology and evolution

In order to address questions in infectious disease evolution, we developed a library for flexible epidemiological simulations of evolving pathogens. The library is named Opqua, after the word for disease, cause, or reason in the Chibcha family of languages spoken by members of the late Muysca Confederation in modern-day Colombia (*47*).

Opqua stochastically simulates interconnected populations of agents representing hosts and/or vectors, which may be infected with pathogens (Fig. 1). Simulations occur through different kinds of demographic, epidemiological, immunological, and genomic events that affect the state of the system, and thus the probability of the next event (Fig. 1B; Fig. S1). User-defined epidemiological parameters affect the rate of each event and can vary both across populations and throughout the course of each simulation, allowing the user to model different ecological events or public health interventions (Fig. 1C). Hosts and vectors may acquire immunity to specific pathogen genome sequences, and may undergo demographic change by natural death and reproduction, as well as migration within complex population structures (Fig. 1D).

**Fig. 1.**
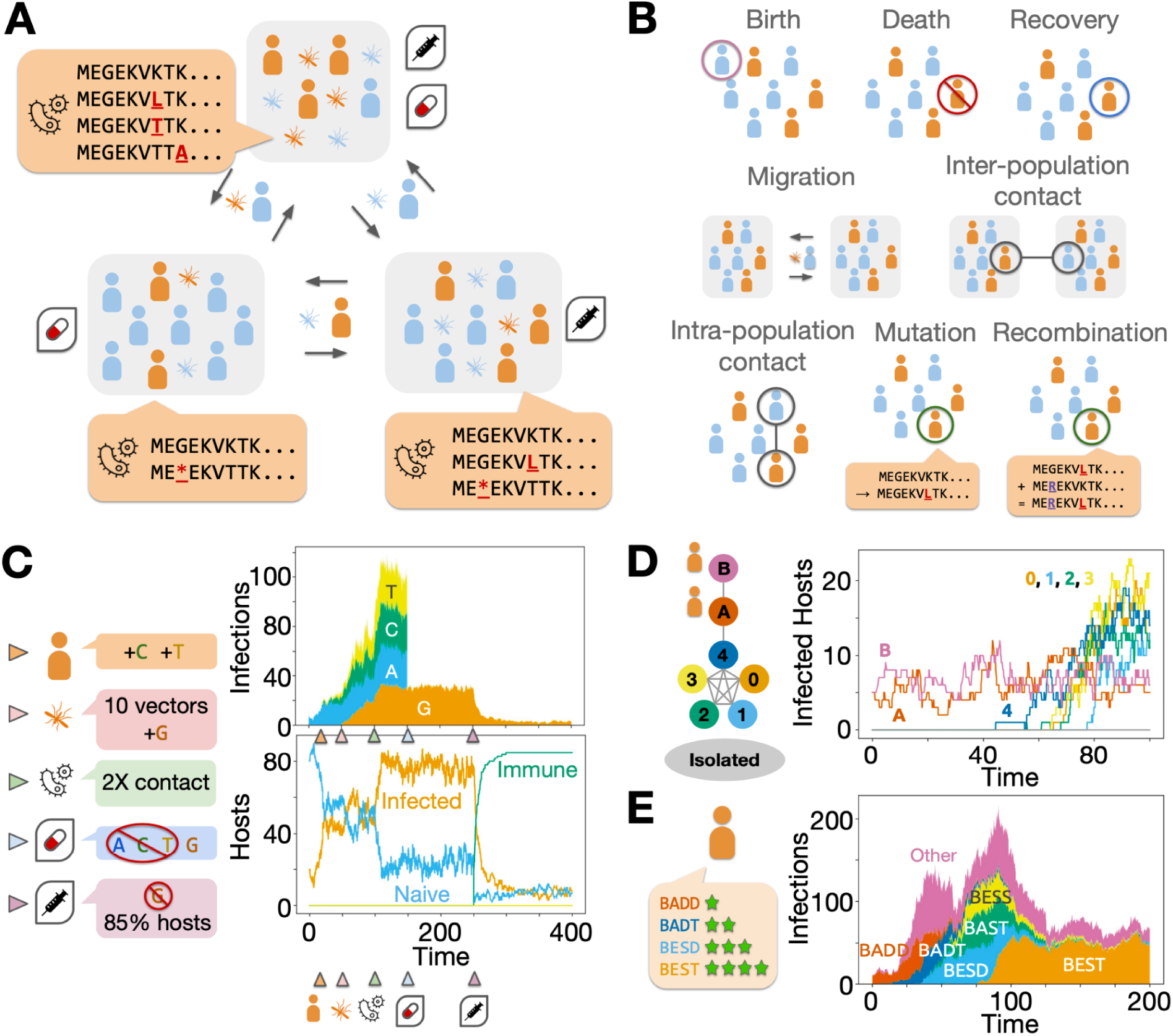
Opqua simulates stochastic, structured epidemiological models of evolving pathogens. **(A)** Opqua models account for pathogens with arbitrary genomes and genomic alphabets spreading through structured populations of hosts and/or vectors, which may acquire immunity and/or undergo interventions such as drug treatment or vaccination. **(B)** Eight possible kinds of events can occur in simulations, the rates of which may be influenced by pathogen genome sequence. **(C)** Models may include interventions at different moments in time, which in this example include addition of new hosts and vectors infected with pathogens of specific genome sequences, altering epidemiological conditions to increase host-vector contact rate, administration of a treatment that kills all pathogens except those with a specific, resistant genotype, and finally, vaccination against the same genotype. **(D)** Models may have arbitrary, complex metapopulation structures, here showing the spread of disease from two low-transmission populations (“*A*” and “*B*”) into five interconnected populations with high transmission (labeled 1–5), while an additional isolated population remains disease-free. **(E)** Models can simulate evolution and selection through different ways, here showing a population of pathogens that evolve through direct intra-host competition and *de novo* mutation from an initial, low-fitness genotype (“BADD”) to a high-fitness, optimal genotype (“BEST”).

Pathogens can be transmitted through direct contact among or between hosts and/or vectors, or through vertical transmission. Pathogens have genomes composed of segments or chromosomes represented by sequences of letters from an arbitrary alphabet, such as DNA bases, amino acids, or other arbitrary alleles. Genomes are capable of mutation, chromosome reassortment, and recombination. Crucially, genome sequences may affect the behavior of their associated pathogens and the hosts and vectors they infect by modifying the rates of any event, resulting in selection and complex evolutionary dynamics (Fig. 1E). Finally, Opqua contains functions to graph and analyze the evolutionary and epidemiological trajectories of the simulation. The software is available as a Python package with documentation, examples, and source code (github.com/pablocarderam/opqua), and a detailed discussion of its functioning in the supplementary materials.

### Effect of transmission environment on evolution across fitness valleys

Using Opqua’s evolutionary modeling capabilities, we examined the epidemiological determinants of pathogen stochastic tunneling across fitness valleys (*39, 41*). We began by examining whether competition for transmission between pathogens in high transmission environments can inhibit stochastic tunneling. This hypothesis has been proposed as a partial explanation for why antimalarial resistance repeatedly emerges outside of the high-transmission environments in sub-Saharan Africa that contribute the bulk of global malaria cases and treatment (*37, 48, 49*).

We established a simple model in which mutating pathogens compete with each other for transmission between hosts within a single population. Pathogen fitness follows a valley-like pattern in which an initial fitness peak of “wild-type” genomes is separated by a sequence of mutations from a second, higher fitness peak of “resistant” genomes capable of surviving drug treatment and causing longer infections (Fig. 2A). When pathogens with different genomes coexist within the same individual, those with more fit genomes have a greater probability of transmission and mutation, due to their greater share of the intra-host pathogen population. After letting the system evolve for a certain amount of time, we apply a drug treatment that kills all pathogens, save for those with the resistant genome.

**Fig. 2.**
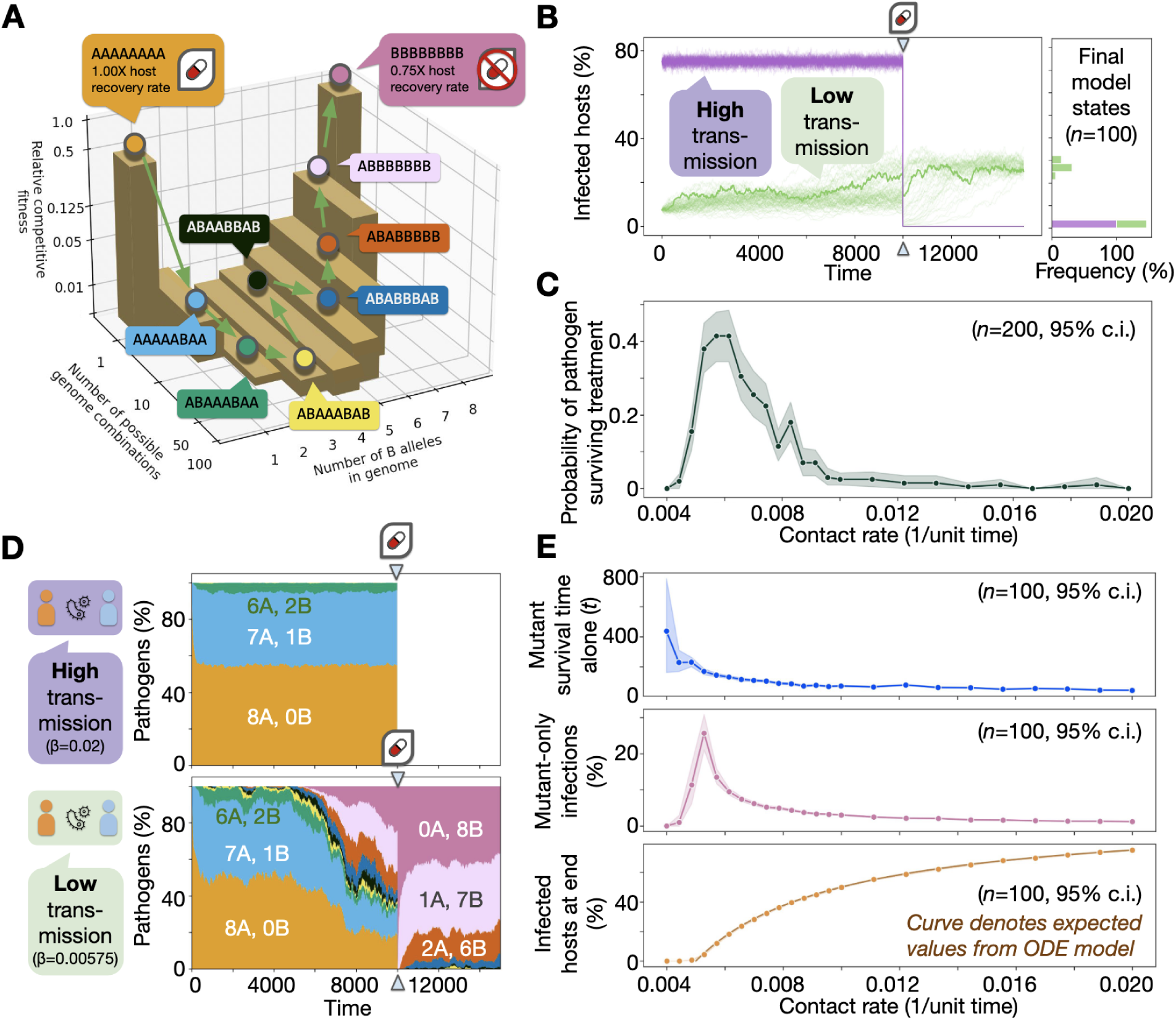
Competition between pathogens restricts evolution across fitness valleys. **(A)** A model system contains pathogen genomes with eight different bi-allelic loci. Fitness follows a valley-like distribution according to a genome’s “B” alleles, where the “0-B” wild-type sequence outcompetes all mutant genomes except for the “8-B” sequence, which encodes for higher competitive fitness, lower recovery rates, and drug resistance. **(B)** Over time, tracking infections in 1000 hosts shows that pathogens in a high-transmission environment are eradicated during a drug administration event. However, drug effectiveness is halved in a low transmission environment, as resistant mutants can evolve. **(C)** An optimal, low contact rate favors the evolution of the drug resistance trait in this system. **(D)** By tracking the genetic composition of the pathogen population over time, we see that at high contact rates (*β*), most mutants remain evolutionarily close to the wild-type, whereas low contact rates lead to the breakthrough and population sweep of distant mutants with high fitness. **(E)** Even when removing all mutant advantages, the frequency and duration of mutant-only host infections is greater at low contact rates, but is counteracted by low infected host totals. The interplay between these two forces determines stochastic tunneling across the fitness valley.

By simulating disease outbreaks in this model system, we are able to recapitulate the predicted competitive interference behavior (Figs. 2B, 2C). At high transmission intensities, pathogens fail to evolve resistant genotypes and the infection is cleared from the population after drug treatment. However, low transmission settings allow mutant pathogens to persist and evolve across the fitness valley and reach the more fit, resistant genotype, which is able to survive drug treatment.

We can track the genomic composition of populations over time to observe the evolutionary paths taken in each environment (Fig. 2D). When transmission rates are high, mutant pathogens are rarely able to survive for long within hosts without wild-type pathogens coinfecting and outcompeting them. In contrast, low transmission environments allow mutants to evolve for longer without interference from wild-type pathogens. This leads to a greater frequency of distant mutants in low transmission environments, which eventually result in the appearance and fixation of resistant pathogens. Thus, low transmission environments generate the cryptic genetic variation that has been shown to underlie evolution to new adaptive peaks in other experimental and theoretical models (*50, 51*).

However, by varying the contact rate, we can see that the relationship between transmission and evolution is not linear (Fig. 2C). Instead, the likelihood of pathogens evolving resistant genotypes is a result of two opposing forces. On one hand, there is the likelihood of having hosts infected by low-fitness mutants for long periods of time without wild-type competitors that interfere with further evolution towards more distant genotypes. This likelihood is greater in low transmission environments where competition is weak, as long as transmission is strong enough to support a stable pathogen population (Fig. 2E, top and middle). On the other hand, we have the likelihood of survival for strains that reach new fitness peaks. This likelihood is greater in high transmission environments, which avoid stochastic extinction events (Fig. 2E, bottom). The balance between these opposed dynamics determines an optimal transmission level for evolution (Fig. 2C), dependent on the features of the adaptive landscape.

In particular, low transmission regimes can only increase stochastic tunneling through fitness valleys if the mutational distance between peaks and the drop in fitness between them is large enough to require evolution to occur throughout a transmission chain of multiple hosts (Fig. S2). Long chains of transmission facilitate tunneling through long, deep valleys due to multiple bottlenecks and opportunities to minimize pathogen competition. However, shorter valleys not only permit stochastic tunneling to occur in short chains of transmission (or even single hosts), but also imply a more restricted number of mutational paths through the valley. This decreases the likelihood of low transmission regimes leading to stochastic tunneling due to their smaller pathogen populations and thus net mutation rates. In this way, high transmission environments are more effective at taking single evolutionary steps with significant fitness differences, whereas low transmission environments are more effective at taking multiple, successive steps with minor differences in fitness. Our results show these small steps are ultimately more effective at tunneling through long evolutionary distances with low overall fitness.

### Evolution across descending fitness landscapes in deterministic and stochastic models

We next focused on the evolutionary dynamics of pathogens mutating into low fitness regimes. To compare genomic epidemiological models and traditional epidemiological modeling techniques, we constructed a compartment model describing a host-pathogen system in which uninfected hosts can become infected with wild-type and/or mutant genotypes (Figs. 3A, S3). Similar to our previous models, mutant pathogens have lower competitive fitness than their wild-type counterparts.

**Fig. 3.**
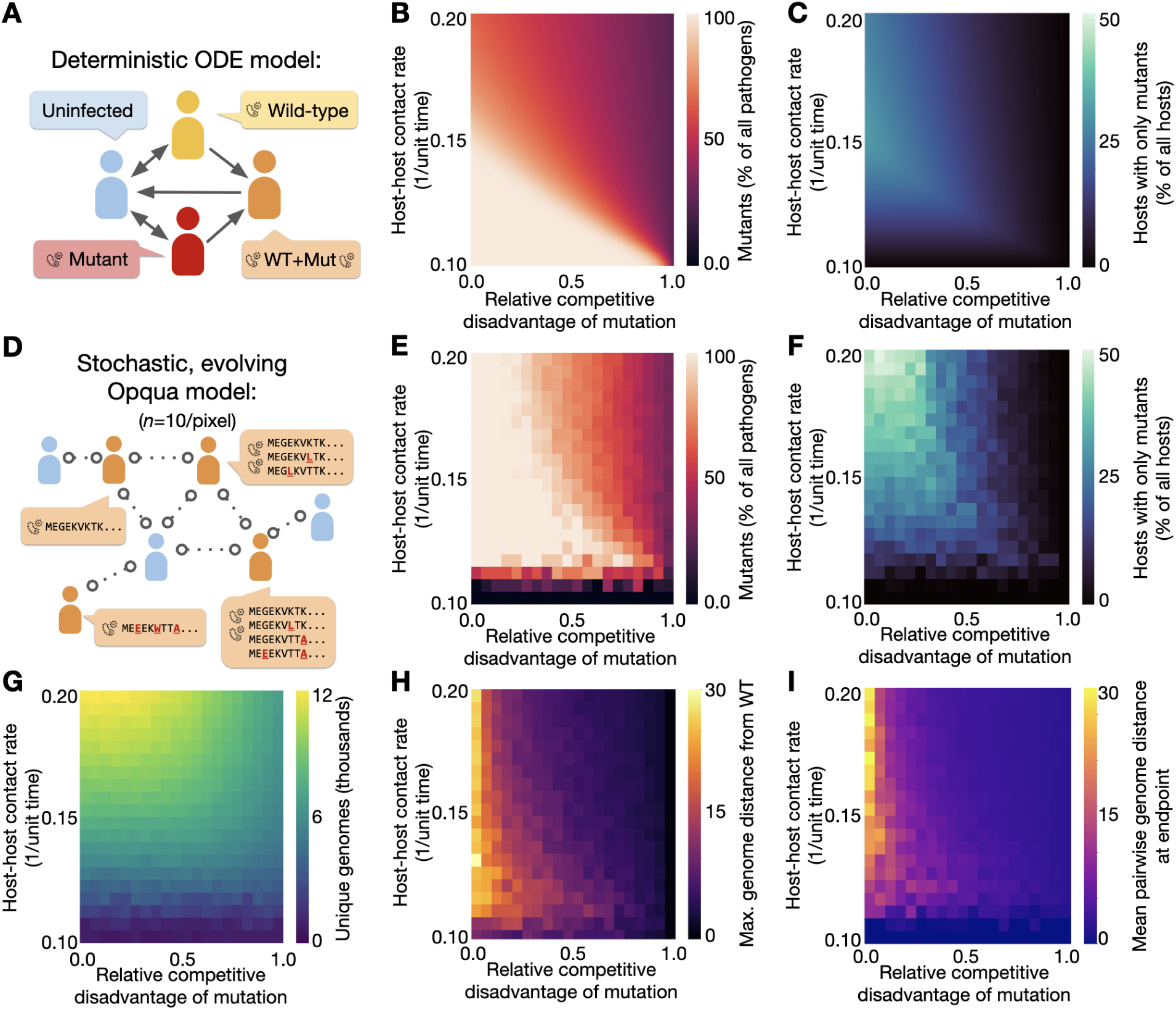
Deterministic and stochastic simulations describe the effect of competition on the distribution and dimensions of pathogen evolution. **(A)** Ordinary differential equations (ODE) model the evolution of a pathogen with a single bi-allelic locus. (**B**) Mutant pathogens are still common at high fitness costs to mutants, as long as contact rates are low. (**C**) Lone mutants (with no within-host wild-type competitors) are favored at a peak contact rate, which lowers in value as the fitness cost of mutation rises. (**D**) A stochastic simulation of a similar system tracks evolution across 500 loci, each with 20 possible alleles. Each mutation away from the wild-type sequence reduces fitness by the factor shown. The distributions of mutants among (**E**) pathogens and (**F**) hosts show similar behaviors to the ODE model. Stochastic pathogen extinction events drive mutants down at low contact rates, while genetic drift drives them up at high contact rates. Furthermore, simulations allow analysis of the resulting genomes. (**G**) High contact rates always lead to a higher number of genotypes explored. However, when mutation fitness costs increase, low contact rates allow for the evolution of pathogen lineages more distinct both from (**H**) the wild-type and (**I**) each other.

We numerically solved this model while varying the relative competitive disadvantage of mutants and the intensity of transmission. The results are concordant with our earlier findings, showing that low transmission environments facilitate the survival of unfit mutants in a pathogen population, even when the fitness cost of mutations is considerably high (Fig. 3B). The model also shows an optimal transmission intensity at which “mutant-only” infections peak for any given mutation fitness cost (Fig. 3C), similar to the results of the fitness valley model. The optimal transmission level for mutant-only infections decreases as the competitive fitness disadvantage of mutants becomes more prominent. We also varied the infected host fraction while keeping contact rates constant by changing the host recovery rates, which showed essentially the same dynamics of mutant prevalence as seen from the contact rates studied (Fig. S4). This shows that the differences in mutant prevalence at different contact rates arise mainly from competition between wild-type and mutant pathogens.

We then constructed a stochastic model using Opqua to study the same type of behavior in a genomic context. This model had epidemiological parameters identical to those in the compartment model, with the only difference of allowing hosts to become infected with pathogens containing complex genomes simulating a peptide sequence of interest with 500 amino acids (Fig. 3D). Each mutation decreased pathogen fitness by a set factor, resulting in an exponentially-decreasing fitness landscape away from the single peak of the wild-type genome sequence. This descending fitness regime, analogous to an infinitely long valley, can be used to simplify the study of evolution in the initial part of a fitness valley.

By plotting the average of multiple simulation replicates at different contact rates and mutant fitness costs, we reproduced the behaviors observed in the compartment model, with two differences (Fig. 3E, 3F). First, contact rates that are too low in the genomic Opqua model result in stochastic extinction of all pathogens (counted as a zero fraction of mutants). Second, the genomic model shows greater mutant prevalence at low mutation fitness costs and higher contact rates than the equivalent parameters in the compartment model. This is a consequence of genetic drift: if different genomes have comparable fitness and transmission is not a scarce resource, the wild-type genome becomes just one among a large number of similar, competing genotypes, and is subject to stochastic extinction like its peers.

Furthermore, genomic models allow us to analyze the sequences and distributions of genomes that evolved throughout the simulations and construct a quantitative description of the different kinds of evolutionary space explored. For instance, high transmission regimes always result in a higher number of genomes explored, due to higher numbers of infections providing a greater net mutation frequency (Fig. 3G). However, even though low contact rates result in fewer mutants, those mutants can evolve further from the wild-type starting point even at high fitness costs to mutation (Fig. 3H), echoing the results of the fitness valley model. The resulting genomes at the end of the simulation are also more different from each other at low contact rates (Fig. 3I). Thus, genomic epidemiological models more accurately capture the behavior of systems with high genetic dimensionality and provide a quantitative description of how transmission environment and within-host competition shape pathogen cryptic variation.

### Effects of host population structure and pathogen biology on evolution

With these metrics of genomic evolution, we examined other kinds of epidemiological determinants of evolution beyond transmission intensity, starting with host population size and inter-population mobility (Fig. 4).

**Fig. 4.**
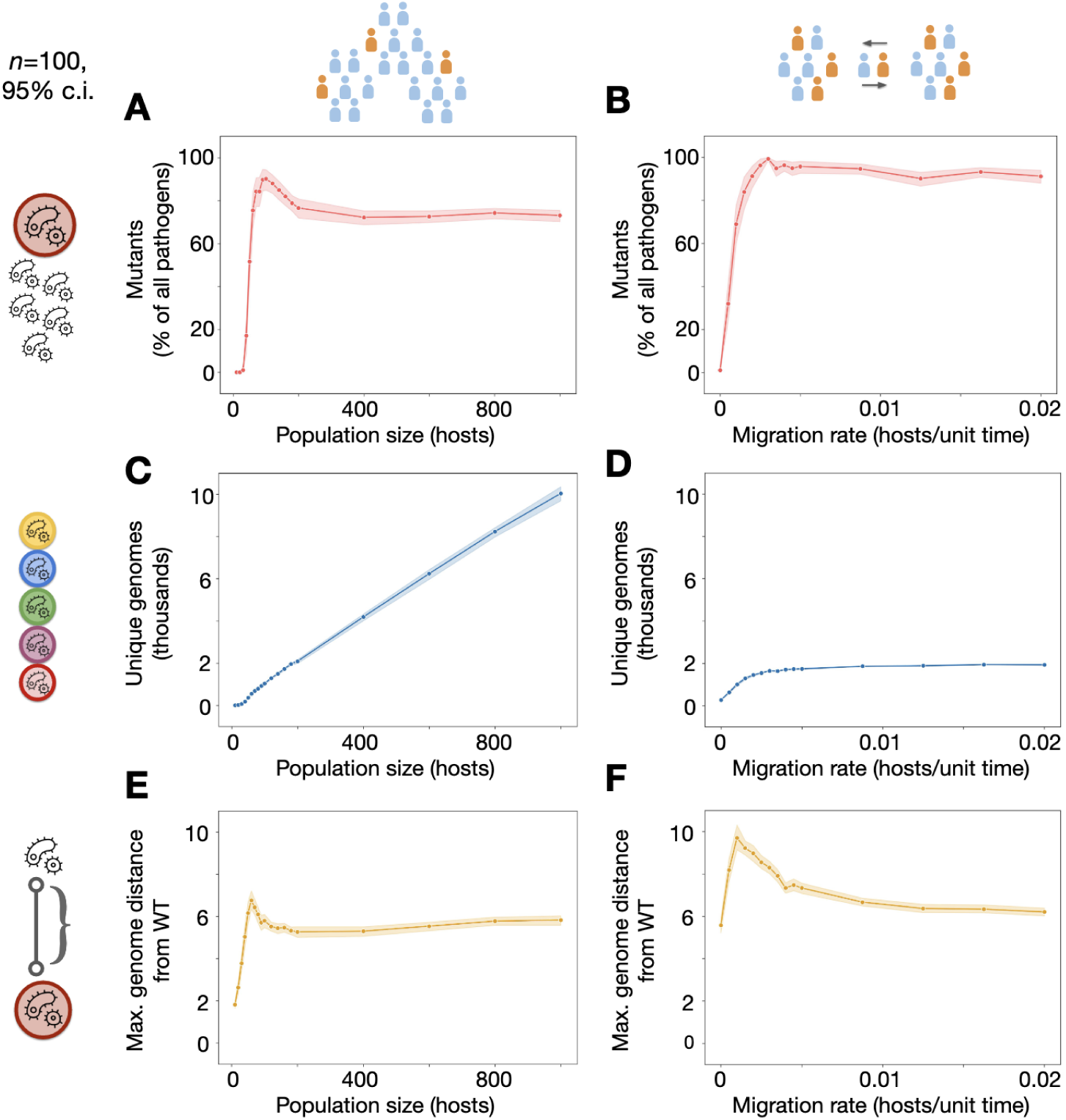
Population size and structure affect the distribution and dimensions of pathogen evolution. **(A)** At constant contact rates and relative competitive fitness of mutants, population sizes that are too low cause stochastic extinction of mutants. Mutants increase in frequency as host populations grow, aided by fixation through genetic drift. At larger population sizes, drift leading to wild-type extinction is less likely and the mutant population stabilizes at a lower value. **(B)** The frequency of mutants within the pathogen population follows a similar pattern when increasing inter-population host migration in a system of 10 interconnected sub-populations, each initially with 10 hosts. **(C)** An increase in host population size leads to a near-proportional increase in the number of unique pathogen genomes explored. **(D)** In contrast, increasing mobility leads to the number of genomes explored converging on the value for a single population of 100 hosts. **(E)** Genome distance from the wild-type sequence follows a similar pattern to mutant fraction when increasing population size. **(F)** Low migration rates with frequent local extinction and re-colonization events allow pathogens to evolve across longer evolutionary distances than what either the individual population or total metapopulation sizes (10 and 100 hosts, respectively) would normally permit.

Increasing population sizes initially leads to a greater fraction of mutants among the pathogen population, since small host populations lead to stochastic extinction events for all pathogens (including mutants, which is counted as zero; Fig. 4A). At slightly larger population sizes, the pathogen population is stable enough to avoid complete extinction, but small enough to be susceptible to genetic drift and allow slightly unfit mutants to fixate, driving wild-type pathogens extinct. At even larger host population sizes, the pathogen population is less susceptible to genetic drift and the stochastic extinction of more fit wild-type pathogens, leading to a slightly diminished equilibrium fraction of mutants. We see similar dynamics by varying host inter-population migration rates in a metapopulation model with a fixed number of total hosts (Fig. 4B). Low mobility rates make the metapopulation system behave like a series of small, isolated populations, while high mobility makes the system behave like a single, well-mixed, large population.

However, different behaviors emerge when examining genomes. While larger populations lead to more infections and thus a greater number of unique genomes explored (Fig. 4C), increased mobility does not increase infected hosts, and therefore unique genomes, beyond a certain point (Fig. 4D). The stochastic extinction of high-fitness wild-type pathogens in small populations allows for an early peak in genome distance from wild type, before descending to a slightly lower stable value for larger populations (Fig. 4E). Intriguingly, the most notable increase in the maximum evolutionary distance explored comes from low host mobility systems, provided host mobility is high enough to sustain continued infection (Fig. 4F). Low mobility between host populations results in local pathogen extinctions that allow any mutant pathogens arriving in the population to spread without competition. In this way, the reduced genetic variability initially produced by founder effects paradoxically generates increased variability in the long run. This result is analogous to the effects of low transmission within populations shown in our previous models, with similar evolutionary mechanisms operating on a different scale.

Finally, we used the same approach to examine how different aspects of pathogen biology shape evolution in the kinds of descending fitness regimes explored earlier. We focused on the effects of *de novo* mutation rate, the number of pathogens inoculated during infection, and genomic recombination rates in two modes of transmission: host-host (Figs. S5, S6) and host-vector. Transmission in the host-vector model was accompanied by selection in hosts only (Fig. 5), or separate selection regimes within hosts and vectors (Figs. S7, S8).

**Fig. 5.**
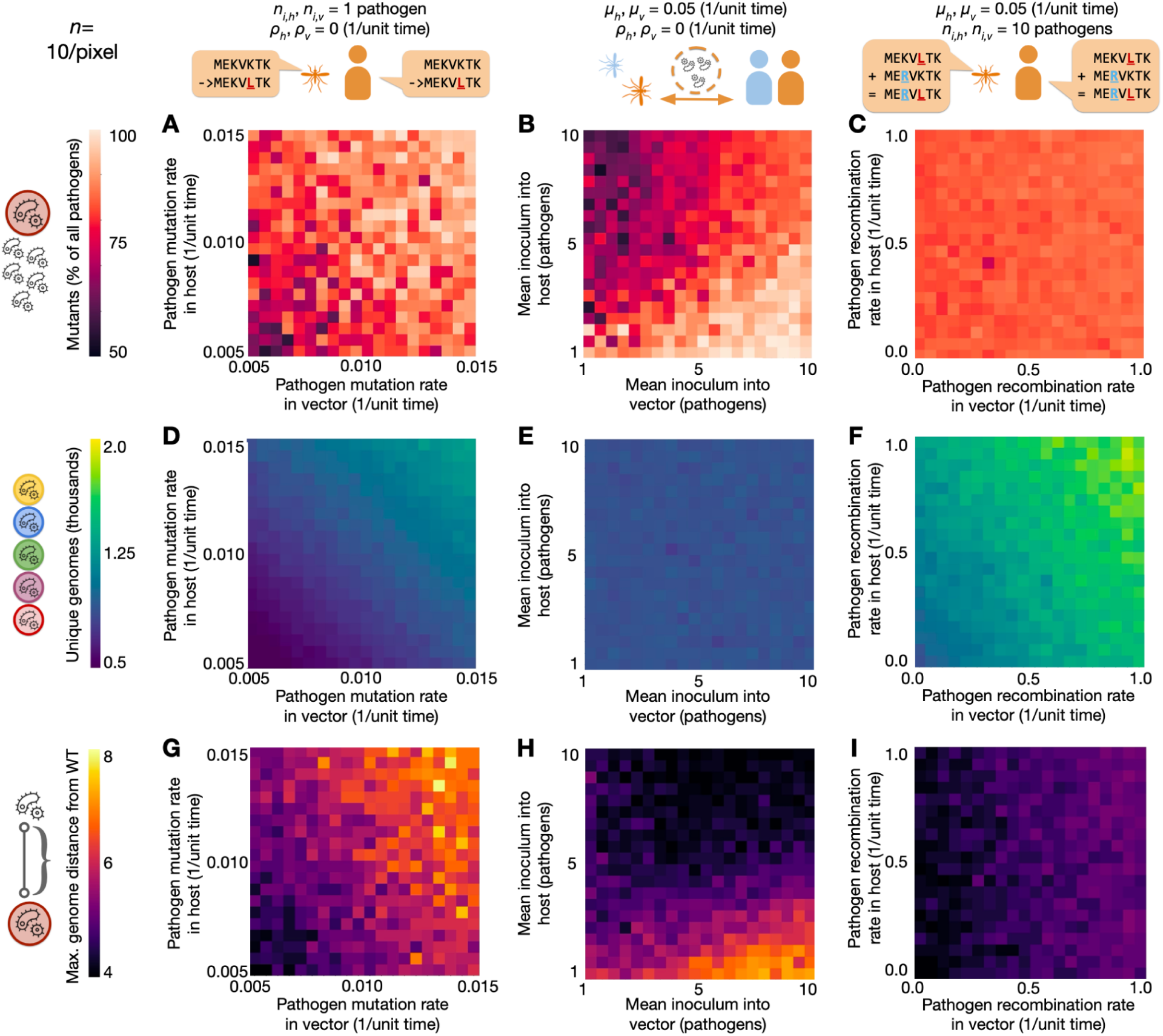
Pathogen biology and life history affect the distribution and dimensions of evolution. **(A)** High mutation rates in both hosts (*µ_h_*) and vectors (*µ_v_*) lead to greater mutant frequency among pathogens. **(B)** Greater mean inoculum sizes from hosts (where competitive selection is present) into vectors (*n_v_*) increase mutant frequency by capturing within-host diversity. However, mutant frequency increases at lower inoculum sizes from vectors (where selection is absent) into hosts (*n_h_*), due to bottlenecks reducing intra-host competition. **(C)** Recombination rates have little effect on mutant frequency. **(D)** Increasing mutation rates in both hosts and vectors leads to a higher number of genomes explored, while **(E)** inoculum sizes into both hosts and vectors have no effect. **(F)** Increased recombination rates in both hosts and vectors lead to an increase in the number of genotypes explored. **(G)** The maximum evolutionary distance from wild-type (WT) increases with mutation rates. **(H)** Low inoculums into hosts and high inoculums into vectors are optimal for increasing evolutionary distance, as they combine bottlenecks before selection and capture diversity after it. **(I)** Increased recombination within vectors (where competitive selection is absent; *ρ_v_*), more so than hosts (where competitive selection is present; *ρ_h_*), leads to higher distances from wild-type.

Vector-borne pathogens often have vastly different genetic programs and host-pathogen interactions in different life stages, leading to an uncoupling of selection in which mutations affect pathogen fitness differently in hosts and vectors. In some cases, this can lead to selection only occurring within one kind of organism (such as the main hosts), while all genomes have equal competitive fitness within the other (such as the vector organisms) (Fig. 5). As an example of this, it is well known that multiple genes in *Plasmodium* malaria parasites are expressed only in human red blood cell-infecting stages and not mosquito-infecting stages, or vice-versa (*52–54*). In repeated instances, parasite genes knocked out with no detectable phenotype in one set of life stages are found to be essential for survival in other life stages (*55, 56*). Nevertheless, other situations can lead to a different kind of uncoupling, where selection is present in both hosts and vectors, but follows two different fitness landscapes (Figs. S7, S8). This may occur, for instance, if a gene fulfills roles in host-pathogen interactions or is subject to different kinds of immune responses within both hosts and vectors, as is common in many arboviruses (*57, 58*). It may also be a consequence of antimicrobial use if a resistant allele with fitness costs is selected for in hosts (but not vectors) due to drug treatment (*45, 46*).

Unsurprisingly, increasing mutation rates always increase mutant prevalence among pathogens (Figs. 5A, S5A, S8A), the number of unique genomes (Figs. 5D, S5D, S8D), and the maximum distance from wild-type (Figs. 5G, S5G, S8G). However, varying inoculum size from hosts into vectors and vice-versa in a host-vector system shows complex interactions between the two variables (Fig. 5H). Consider a regime of host selection only (Fig. 5H) and small inoculums into hosts. In this setting, small inoculum sizes into hosts are key to increasing mutant fractions of the pathogen population (Figs. 5B, S5B) and maximum evolutionary distance from wild-type (Figs. 5H, S5H). This is due to the fact that small inoculums create a population bottleneck that allows mutants to be transmitted without wild-type competitors. Additionally, when inoculating from vectors into hosts, pathogens are sampled randomly without a transmission disadvantage for mutants. Combining small inoculums from vectors into hosts with a high inoculum into vectors captures greater genetic variation generated in the host, leading to maximized mutant fractions and evolutionary distance from wild-type (Fig. 5H). Paradoxically, if the mean inoculum into hosts is high, the effect is reversed. In these cases, contacts from infected vectors are more likely to result in coinfections in hosts, leading to competitive exclusion of unfit mutants. Therefore, when inoculums into hosts are high, small inoculums into vectors reestablish the population bottleneck that reduces competition from wild-type pathogens, leading to greater mutant fractions and evolutionary distance from wild-type.

The effect of these processes is replicated symmetrically when considering selection regimes in both hosts and vectors (Fig. S8H). In this case, the greatest long-distance evolution occurs when a mismatch in inoculum sizes is present (this is to say, a high inoculum from hosts to vectors combined with a low inoculum from vectors to host, or vice-versa). Since there are selection pressures in both life stages, bottlenecks reducing competition and amplifications capturing variation enhance evolution in either stage. However, if inoculums into both hosts and vectors are large, competition between pathogens suppresses long-distance evolution. Conversely, small inoculums in both directions capture less genetic variability. This also decreases evolution, although to a lesser extent. It is worth pointing out that the dynamics of evolutionary distance (Fig. S8H) and mutant pathogen populations (Fig. S8B) become uncoupled in this case, due to the opposing effects that bottlenecks in transmission to each life stage have on mutant fractions.

Recombination presents another interesting case where the dynamics of mutant pathogen fractions and maximum evolutionary distance become uncoupled. Because recombination cannot create *de novo* genetic variation, mutation rates still largely determine mutant prevalence. This means the effect of recombination on mutant fractions of pathogen populations is minimal (Figs. 5C, S5C, S6A, S8C). Nevertheless, recombination rates in both hosts and vectors increase the number of unique genomes explored (Figs. 5F, S5F, S6B, S8F). Additionally, recombination can increase the maximum evolutionary distance from wild-type explored by pathogens. In a host-vector system with selection in hosts only, the effect of recombination on evolutionary distance is largest if recombination occurs within vectors, a life stage where the new genotypes will not be outcompeted by their parents (Fig. 5I). If selection occurs with separate fitness landscapes in hosts and vectors, the effect becomes symmetrical for both life stages, due to recombination of genomes clustered around each of the two peaks (Fig. S8I). Nevertheless, recombination in hosts can also increase the maximum evolutionary distance in host-host models (Fig. S5I). The magnitude of this effect is dependent on the mean inoculum size (the infection effective population size), as small inoculums remove the within-host variation from which recombination can occur (Fig. S6). This shows recombination can also play a relevant role in stochastic tunneling of infections with host-host transmission.

### Perspectives and outlook

In this work, we present Opqua, a flexible computational modeling framework capable of simulating genomic epidemiology. We use it to study pathogen competition and evolutionary dynamics in fitness valleys and descending fitness regimes with a quantitative, systematic approach. This allows us to better understand how pathogens acquire new fitness-conferring traits that require multiple, separate epistatic mutations, a process known as stochastic tunneling (*39, 41*). We confirm that competition between pathogens in settings with high disease prevalence can pose a significant barrier to evolution in certain adaptive landscapes, as suggested by others (*37*). We establish the potential of genomic epidemiological models to provide more complete descriptions of simulated evolution. We then use these descriptions to examine how different aspects of host populations and pathogen biology can alter the competitive and genetic population dynamics that shape evolution. By simulating these traditional scenarios from theoretical evolutionary biology in an epidemiological context, we provide concrete illustrations of counterintuitive phenomena that can aid decision making in disease control. Specifically, we show that stochastic effects in small, relatively isolated, structured populations can increase the evolutionary distance traversed by pathogens, exploring evolutionary space in greater depth. Lastly, we show how decoupling selection from transmission in pathogen life cycles with multiple stages changes the evolutionary dynamics across stretches of low fitness. Mismatches in inoculum sizes during transmission from one life stage to another and genetic recombination in changing selective regimes increase long distance evolution.

These findings suggest distinct strategies with evolutionary advantages. For pathogens that depend on direct contact between hosts, such as viral respiratory infections, inoculum sizes are often inextricably tied to transmissibility, resulting in a trade-off between transmission and stochastic tunneling to new adaptive peaks. Conversely, infections with complex life cycles can randomly subsample genetic variability within life stages where selective pressure is weak to increase the maximum evolutionary distance explored, leading to more stochastic tunneling. This can be seen in malaria parasites, which undergo strong population bottlenecks in mosquitoes and within human livers, before intra-host competition in the bloodstream occurs. These bottlenecks reduce the number of individual pathogens by ten orders of magnitude from its highest point within the human bloodstream (*59*), and have been shown to qualitatively alter the pathogen’s evolutionary dynamics (*60*). Our findings corroborate this and indicate that interventions that target these vector stages and their bottlenecks not only reduce disease prevalence, but can inhibit pathogen evolvability if they introduce selective pressures on traits that affect host stages.

Our results also have important eco-epidemiological implications. When considering the evolution of distant pathogen genotypes with higher fitness through stochastic tunneling, rural communities consisting of small, semi-isolated groups with relatively low transmission may be more at risk than larger populations with high transmission. It is interesting to consider that Omicron’s emergence may well be an example of this kind of stochastic tunneling in small, remote populations with cryptic transmission, as proposed by others (*61*). Other explanations such as evolution in individual chronic infections or through reverse zoonosis have found some evidence in their favor as explanations for the emergence of Omicron, and it is doubtless that selective pressure from acquired immunity played a key role (*62*). However, it is worth noting that our results show that no special population compartments (such as chronically infected individuals or other animals) are needed to explain the fact that the chain of mutations that led to Omicron went undetected for over a year, the time separating the last common ancestor between Omicron and other sequenced variants [nextstrain.org/groups/neherlab/ncov/21K.Omicron, (*22, 63*)]. Stochastic tunneling is just more likely to occur in remote populations outside of surveillance efforts. Exploring the likelihood of this hypothesis with respect to others in the emergence of Omicron is beyond the scope of this work. Regardless, our results suggest that novel, evolutionary distant variants are more likely to emerge in the places where detection would be most challenging: fragmented, mobile populations in regions with relatively low incidence and cryptic transmission of low-fitness genotypes. In this light, monitoring pathogen genomes in peripheral communities could be of great value for genomic surveillance programs. This runs contrary to conventional practice in national COVID-19 surveillance programs, which has so far focused on monitoring large, central population hubs (*9–15*).

A final corollary of these results is that under certain conditions, pathogen evolvability increases the more elimination efforts progress against it. The low-transmission environments brought about by eradication campaigns are the exact conditions needed for stochastic tunneling. This does not in any way imply that eradication efforts are pointless—after all, evolution is always a possibility, while disease burden is a certainty. Nevertheless, it might help explain why elimination campaigns are so challenging to complete, despite initial success.

We deliberately chose to focus on intra-host competition, as this is the minimum requirement for blocking stochastic tunneling through competitive exclusion. Therefore, we avoided using Opqua’s ability to modify the intrinsic transmissibility of pathogens according to genome sequence, even though this would increase the magnitude of the effects studied. In addition, by choosing to center on competitive dynamics, we opted to keep the effects of acquired immunity beyond the scope of this study, even though Opqua has the capabilities to simulate them. However, immune selection plays a central role in pathogen evolution by shifting the fitness landscape over time. This has been proposed in the context of the evolution of malaria parasites (*37*) and is doubtless a crucial factor in the emergence of SARS-CoV-2 variants that evade acquired immune responses (*62*). Finally, we chose to explore idealized disease models rather than fit a specific disease both for generalizability and computational feasibility. Future work improving the computational performance of these kinds of modeling approaches may allow for models that more closely fit real-world epidemiological and genetic data, as well as models that simulate within-host dynamics such as drift in a more explicit fashion. Nevertheless, we hope this work helps establish the potential of genomic epidemiological models as a tool to test scenarios, explore hypotheses, and understand the relationship between pathogen evolution and epidemiology. Quantitative descriptions of how biology and environment shape pathogen evolution, such as those presented here, may one day help inform the design of interventions in disease control, ranging from drug and vaccine development to public health policy.

## Materials and Methods

To address the questions posed in this study, we developed Opqua, an epidemiological modeling framework for pathogen population genetics and evolution. Opqua stochastically simulates pathogens with distinct, evolving genotypes that spread through host populations with specific acquired immune profiles. Opqua is available for download through PyPI (pypi.org/project/opqua) with installation, usage instructions, and source code published on GitHub (github.com/pablocarderam/opqua).

### Basic model concepts

Opqua models are composed of populations containing hosts and/or vectors, which themselves may be infected by a number of pathogens with different genomes.

A genome is represented as a string of characters. All genomes must be of the same length (a set number of loci), and each position within the genome can have one of many different characters specified by the user (corresponding to different alleles). Different loci in the genome may have different possible alleles available to them. Genomes may be composed of separate chromosomes or genome segments separated by the "/" character, which is reserved for this purpose.

Each population may have its own unique parameters dictating the events that happen inside of it, including how pathogens are spread between its hosts and vectors.

### Model events

There are different kinds of events that may occur to hosts and vectors in a population:

- contact between an infectious host/vector and another host/vector in the same population (intra-population contact, or “contact”) or in a different population (inter-population contact, or “population contact”)
- migration of a host/vector from one population to another
- recovery of an infected host/vector
- birth of a new host/vector from an existing host/vector
- death of a host/vector due to pathogen infection or by “natural” causes
- mutation of a pathogen in an infected host/vector
- recombination of two pathogens in an infected host/vector

The likelihood of each event occurring is determined by the population’s parameters (explained in the online documentation) and the number of infected and healthy hosts and/or vectors in the population(s) involved. Crucially, it is also determined by the genome sequences of the pathogens infecting those hosts and vectors. The user may specify arbitrary functions to evaluate how a genome sequence affects any of the above kinds of rates. As an example, a specific genome sequence may result in increased transmission within populations but decreased migration of infected hosts, or increased mutation rates. These custom functions may be different across populations, resulting in different adaptive landscapes within different populations.

Contacts within and between populations may happen by any combination of host-host, host-vector, and/or vector-host routes, depending on the populations’ parameters. When a contact occurs, each pathogen genome present in the infecting host/vector is transferred to the receiving host/vector as long as one “infectious unit” is inoculated. The number of infectious units inoculated is randomly distributed based on a Poisson probability distribution. The mean of this distribution is set by the receiving host/vector’s population parameters. The genomes chosen for transmission are sampled randomly according to the fraction of intra-host fitness each genome contributes to the total within the host or vector.

Inter-population contacts occur via the same mechanism as intra-population contacts, with the distinction that the two populations must be linked in a compatible way. For example, if a vector-borne model with two separate populations allows vectors from Population A to contact hosts in Population B, then the vector-host population contact rate in Population A and the host-vector population contact rate in Population B must both be greater than zero. Migration of hosts/vectors from one population to another depends on a single rate defining the frequency of vector/host transport events from a given population to another. Therefore, Population A would have a specific migration rate dictating transport to Population B, and Population B would have a separate rate governing transport towards A.

The recovery of an infected host or vector results in all pathogens being removed from the host/vector. Additionally, the host/vector may optionally gain protection from pathogens that contain specific genome sequences present in the genomes of the pathogens it recovered from, representing immune memory. The user may specify a population parameter delimiting the contiguous loci in the genome that are saved on the recovered host/vector as “protection sequences”. Pathogens containing any of the host/vector’s protection sequences will not be able to infect the host/vector.

Births result in a new host/vector that may optionally inherit its parent’s protection sequences. Additionally, a parent may optionally infect its offspring at birth following a Poisson sampling process equivalent to the one described for contact events above. Deaths of existing hosts/vectors can occur both naturally or due to infection mortality. Only deaths due to infection are tracked and recorded in the model’s history.

*De novo* mutation of a pathogen in a given host/vector results in a single locus within a pathogen’s genome being randomly assigned a new allele from the possible alleles at that position. Recombination of two pathogens in a given host/vector creates two new genomes based on the independent segregation of chromosomes (or reassortment of genome segments, as in influenza virus biology) from the two parent genomes. In addition, there may be a Poisson-distributed random number of crossover events between homologous parent chromosomes. Recombination by crossover event will result in all the loci in the chromosome on one side of the crossover event location being inherited from one of the parents, while the remainder of the chromosome is inherited from the other parent. The locations of crossover events are distributed throughout the genome following a uniform random distribution.

### Model interventions

Furthermore, the user may specify changes in model behavior at specific timepoints during the simulation. These changes are known as “interventions”. Interventions can include any kind of manipulation to populations in the model, including:

- adding new populations
- changing a population’s event parameters and adaptive landscape functions
- linking and unlinking populations through migration or inter-population contact
- adding and removing hosts and vectors to a population

Interventions can also include actions that involve specific hosts or vectors in a given population, such as:

- adding pathogens with specific genomes to a host/vector
- removing all protection sequences from some hosts/vectors in a population
- applying a “treatment” in a population that cures some of its hosts/vectors of pathogens
- applying a “vaccine” in a population that protects some of its hosts/vectors from pathogen infection

For these kinds of interventions involving specific pathogens in a population, the user may choose to apply them to a randomly-sampled fraction of hosts/vectors in a population, or to a specific group of individuals. This is useful when simulating consecutive interventions on the same specific group within a population. A single model may contain multiple groups of individuals and the same individual may be a member of multiple different groups. Individuals remain in the same group even if they migrate away from the population they were chosen in.

When a host/vector is given a “treatment”, it removes all pathogens within the host/vector that do not contain a collection of sequence motifs called “resistance sequences”. A treatment may have multiple resistance sequences. A pathogen must contain all of these within its genome in order to avoid being removed. On the other hand, applying a vaccine consists of adding a specific protection sequence to hosts/vectors, which behaves as explained above for recovered hosts/vectors when they acquire immune protection, in models that allow it.

### Model simulation

Models are simulated using an implementation of the Gillespie algorithm in which the rates of different kinds of events across different populations are computed with each population’s parameters and current state, and are then stored in a matrix. In addition, each population has host and vector matrices containing coefficients that represent the contribution of each host and vector, respectively, to the rates in the master model rate matrix (Fig. S1). Each coefficient is dependent on the genomes of the pathogens infecting its corresponding vector or host. In addition, the contribution of each pathogen genome is weighted by the share of total competitive fitness it holds within the host or vector. This occurs under the simplifying assumption that in a coinfection, pathogens with more competitive fitness will have higher intra-host (or intra-vector) shares of the pathogen population.

This approach also assumes that on the timescales of most short coinfections, pathogen within-host dynamics are driven by selection for fitness-affecting mutations or coexistence in the case of neutral or near-neutral mutations. The model does not currently consider within-host genetic drift. The drawbacks of this assumption are mitigated by the fact that the weight contribution of each pathogen genome affects the event rates associated with it. This means the effect of genome sequence on epidemiology is probabilistic, rather than deterministic, and can approximate the behavior of intra-host drift in the total host population as a whole. Nevertheless, incorporating more realistic (yet computationally-intensive) within-host dynamics is an interesting prospect for future development of the method.

All *x* populations in a model have matrices composed of coefficients *c* for all *n* event types and all *y_h_*hosts or *y_v_* vectors in the population. Hosts and vectors are handled in separate matrices. The coefficient *c* for a given event *e* and host *j_h_*(or vector *j_v_*) in population *i* is computed from the intra-host competitive fitness function *φ* and the family of functions *f*, which return a value between 0 and 1 for all *z* pathogen genomes *g* inside the given host *j_h_* or vector *j_v_* at time *t*:

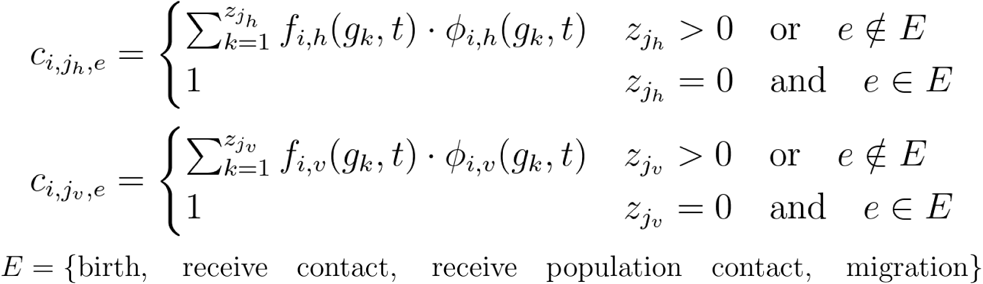

The values of *φ* and *f* are relative to a reference value of 1. Functions *φ* and *f* are defined to be 1 for any given genome *g* by default, but can be set to any arbitrary function that maps a genome sequence to a value between 0 and 1. Each population may have different functions *φ* and *f* for hosts and vectors, and they may be different across populations and time points, as with any other parameter. Coefficients for hosts and vectors with no pathogens are by default zero, save for birth, receiving intra- or inter-population contact, and migration. In these cases, the events may still involve healthy individuals, and the coefficients are therefore equal to 1.

With these coefficients (in units of hosts or vectors for *c_h_* and *c_v_*, respectively), the rates *r* of each kind of event can now be calculated (in units of time^-1^). Some events concern a single host or vector, such as birth, death, mutation, and recombination rates. The total birth rates of hosts and vectors in a population *i* are calculated from the population birth rates *α* (units of time^-1^) at time *t* as

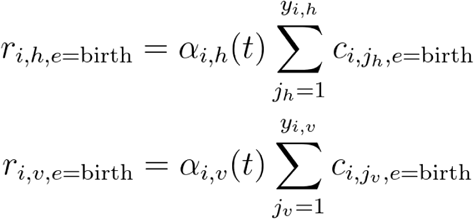

The total natural death rates of hosts and vectors in a population *i* are calculated from the population death rates *γ* (units of time^-1^) at time *t* as

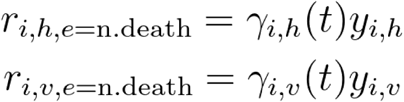

Death by natural causes is unaffected by infection, and thus only depends on the total number of hosts *y_i,h_* or vectors *y_i,v_*. Death due to pathogen infection is counted separately in order to better track the system’s epidemiology. The total mortality rates for hosts and vectors in a population *i* are calculated from the population-specific case mortality rates *τ* (units of time^-1^) at time *t* as

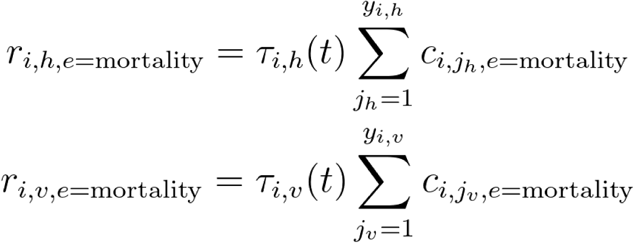

The total recovery rates of hosts and vectors of a population *i* are calculated based on population-specific recovery rates *δ* (units of time^-1^) at time *t* as

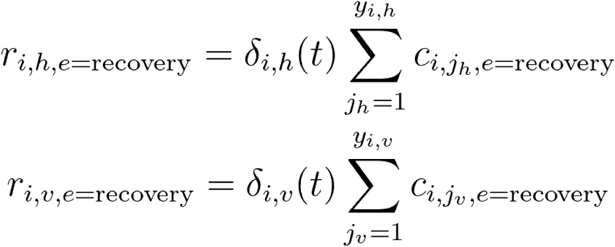

The total rates of *de novo* mutation in hosts and vectors of a population *i* are calculated based on population-specific mutation rates *µ* (units of time^-1^) at time *t* as

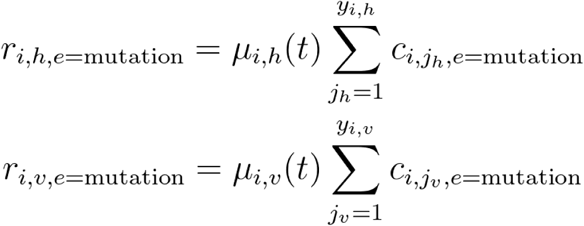

Similarly, the total rates of recombination in hosts and vectors of a population *i* are calculated based on population-specific recombination rates *ρ* (units of time^-1^) at time *t* as

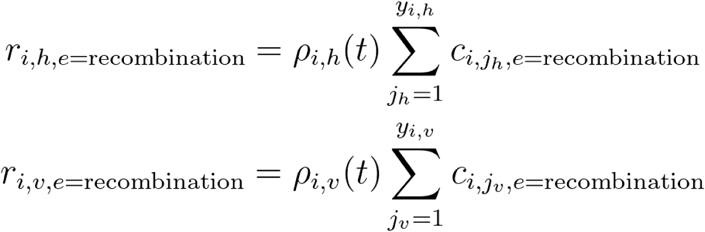

Some events involve two individuals *j_1_*and *j_2_* within the same population. The intra-population contact rates for host-host, host-vector, and vector-host contact in a population *i* are calculated based on population-specific contact rates *β* (units of time^-1^ for host-host transmission, time^-1^host/vector for vector-borne transmission) and transmission efficiency *ɛ* (dimensionless fraction) at time *t* as

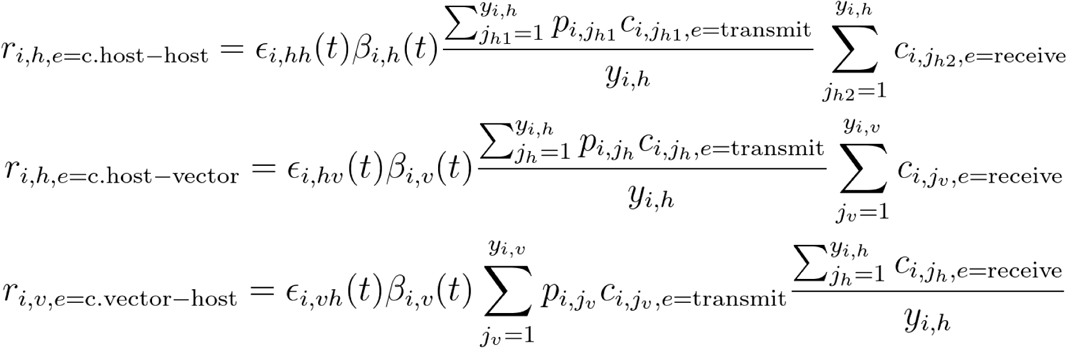

Variable *p* tracks whether a given host or vector is infected with pathogens (*p*=1) or not (*p*=0). Note that for host-vector and vector-host contact, *β* corresponds to *β_v_*and is given in units of time^-1^ host/vector. This follows the convention of defining contact rate in terms of the vector biting rate for parasitic vectors.

Some events involve two different populations. The migration rates of hosts and vectors in a population are calculated based on migration rates *θ* (units of time^-1^) from a specific population *i* to each of all of its *q* possible migration neighbors at time *t* as

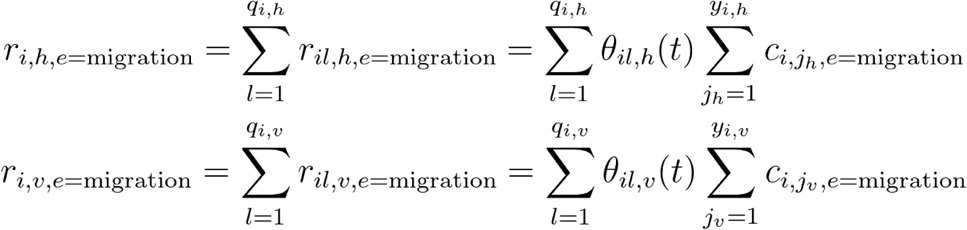

Finally, some events involve two individuals in different populations. The inter-population contact rates for host-host, host-vector, and vector-host contact are calculated based on population-specific contact rates *κ* (units of time^-1^ for host-host transmission, time^-1^host/vector for vector-borne transmission) from a specific population *i* to each of all of its *u* possible contact neighbors at time *t* as

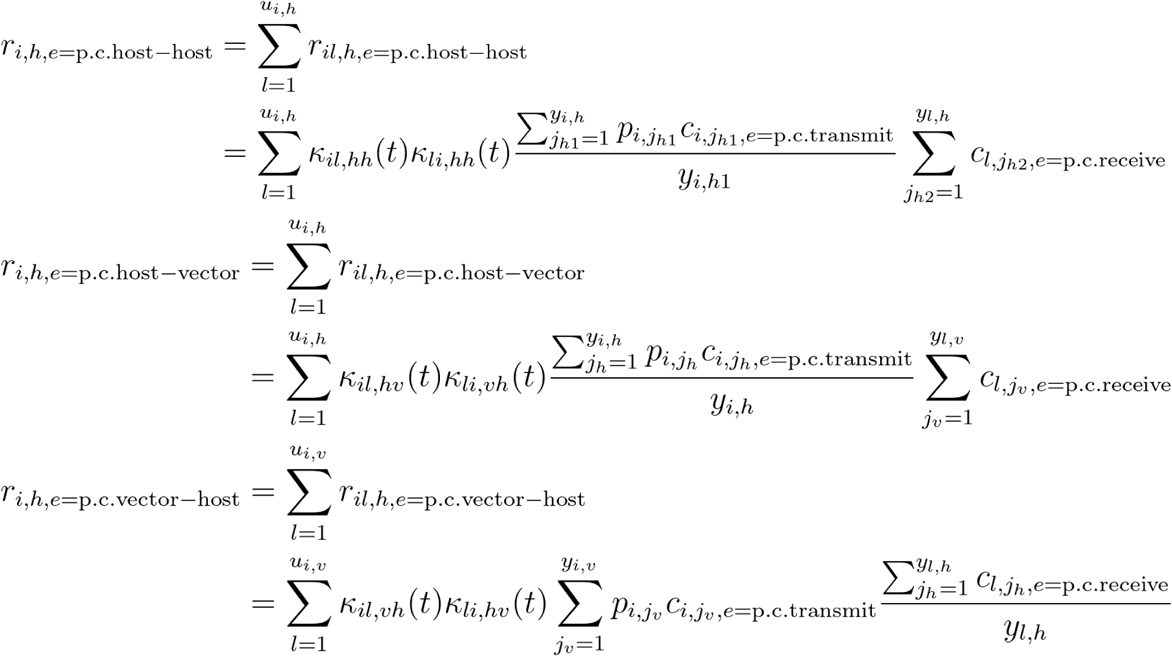

With the rates for all events and populations computed, an event and population is selected randomly, with probabilities drawn from all relative rates as calculated above. In the case of events that involve two populations, the second population is chosen randomly, this time with probabilities drawn from the rates of migration or population contact specifically originating from the first selected population. In similar fashion, the individual host(s) and/or vector(s) are sampled randomly based on the corresponding coefficients within the chosen population(s). Finally, the individual pathogen genomes are sampled randomly based on their respective contributions to the chosen host/vector coefficient. As explained above, mutation events sample a single pathogen, recombination events sample two pathogens, and contact events sample a Poisson-distributed random number of pathogens with replacement. The mean of this distribution is the mean inoculum size parameter, *n_i_*. Whenever an event occurs, the corresponding coefficients for the hosts and vectors affected in the population matrices are updated, and the master rate matrix is recomputed based on this information.

The model’s state at any given time comprises all populations, their hosts and vectors, and the pathogen genomes infecting each of these. A copy of the model’s state is saved at every time point, or at intermittent intervals throughout the course of the simulation. A random sample of hosts and/or vectors may be saved instead of the entire model as a means of reducing memory footprint.

### Model output

The output of a model can be saved in multiple ways. The model state at each saved time point may be output in a single data frame and saved as a tabular file. Other data output types include counts of pathogen genomes or protection sequences for the model, as well as time of first emergence for each pathogen genome and genome distance matrices for every time point sampled.

The user can also create different kinds of plots to visualize the results. These include:

- plots of the number of hosts and/or vectors in different epidemiological compartments (naive, infected, recovered, and dead) across simulation time
- plots of the number of individuals in a compartment for different populations
- plots of the genomic composition of the pathogen population over time
- phylogenies of pathogen genomes

Users can also use the data output formats to make their own custom plots.

### Models used in this study

We used the Opqua modeling framework to study the evolutionary dynamics of pathogens across low fitness regimes. The code to run all simulations and analyses presented in this work is available in a separate GitHub repository (github.com/pablocarderam/fitness_valleys_opqua). The simulations were run on a 2019 MacBook Pro laptop with a 2.4 GHz 8-Core Intel Core i9 processor.

### Stochastic models of fitness valleys

To simulate evolution across a fitness valley (Fig. 2), we constructed a stochastic host-host transmission model using Opqua. The model comprises 1000 hosts in a single population, 500 of which start the simulation infected by “wild-type” pathogens with a genome of “AAAAAAAA”. The epidemiological parameters were set to the default values shown for host-host models in Table S1, with the following modifications: a genome length of 8 loci with two possible alleles (“A” and “B”) at each locus, a mean inoculum of 1 pathogen, a recovery rate of 0.005 1/unit time, a mutation rate of 0.02 1/unit time, and no recombination. The recovery rate of “resistant” pathogens was set to be 75% of the original for pathogens with a genome of “BBBBBBBB”. The choice of using bi-allelic rather than multi-allelic loci (such as four DNA bases, 20 amino acids, or other kinds of allelic variations) was made for simplicity and to more easily observe stochastic tunneling. In systems with many more possible alleles, the probability of reaching a specific other genotype is exponentially lower, and therefore the number of simulation replicates needed to observe differences in stochastic tunneling rates rises exponentially as well.

Most importantly, the intra-host competitive fitness *φ* of each genome was made to follow the following function, describing a fitness valley in terms of the number of “B” alleles in the genome *b* and the total length of the genome *g*:

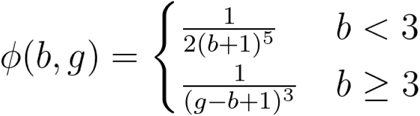

This function was chosen as it describes a steep initial decrease in fitness as pathogens mutate away from the wild type, followed by a more gradual increase in fitness to the resistant genotype. It also guarantees that all mutant pathogens have lower competitive fitness than the wild-type, save for the resistant mutant genome, which outcompetes even the wild type. In this way, resistant pathogens fixate in the population more easily once they arise, which facilitates observation of the stochastic tunneling phenomena being studied.

Additionally, a “drug treatment” intervention was added at time 10,000 such that all pathogens with genomes other than the resistant genotype were cleared from the population.

The simulations shown in Fig. 2E aims to illustrate the evolution of strictly less-fit mutants in the system. Therefore, the previous setup was modified to finish simulation at 10000 time units with no drug intervention, and the contact and mutation rates of resistant pathogens were modified to be equal to zero. This effectively makes the resistant genotype into a lethal mutation, allowing easier study of the remainder of less-fit genotypes without interference. To study the distribution of survival times for mutants that are alone before they become coinfected by a wild-type pathogen or are cleared from the host, we extracted the survival times for 100 mutant-only infections in a single simulation run for each contact rate.

To evaluate the effect of different kinds of fitness valleys (Fig. S2), we varied the number of loci being taken into account for fitness (thus varying the genome length *g* and the length of the fitness valley) as well as *a_d_*, the fraction of the original fitness drop with respect to the wild-type genome (thus varying the minimum fitness of mutants, or the depth of the valley). This was done with the following fitness function:

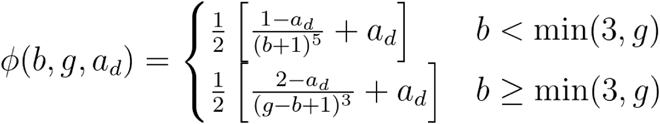

### Compartment model of mutant and wild-type pathogen competition

To study the competitive dynamics of wild-type pathogens and mutant pathogens with lower fitness, we developed a compartment model describing the number of hosts infected with no pathogens (*S*), wild-type pathogens only (*W*), mutant pathogens only (*M*), or coinfected with both kinds of pathogens (*C*). This model is presented in Fig. 3A, and shown in more detail in Fig. S3. The system considers a constant host population *N* such that *N* = *S + M + W + C*. The following system of ordinary differential equations (ODEs) describes the flow between host compartments in terms of host recovery rate *δ*, contact rate *β*, mutation rates from wild-type to mutants *µ_1_* and vice-versa *µ_2_*, inoculum size *n_i_*, and the probability of wild-type pathogens with higher fitness outcompeting mutants in intra-host competition *ψ*:

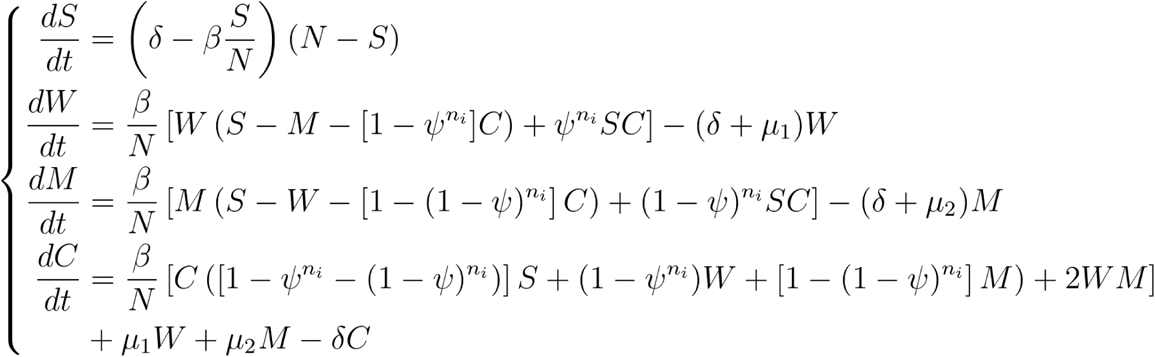

For models in which only a single mutation is possible such as the model above, the relationship between the probability parameter *ψ* used in the ODE model, the relative intra-host competitive fitness *φ* used in Opqua models, and the fitness disadvantage of mutation *λ* shown on Figs. 3 and S4 is given by

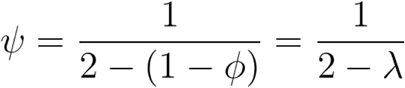

Solving the model at equilibrium leads to *S* = δ N / β* for all nontrivial solutions, but the remaining three steady-state compartments yield unwieldy expressions that are difficult to analyze.

We numerically solved the deterministic behavior of this model using the parameters described in Table S3 from starting conditions of 10 infected individuals, taking the system state after 1000 time units as a result. Mutations that restore the exact wild-type genotype were assumed to be far less likely than mutations away from the genotype (*µ_1_* <<*µ_2_* = 0).

### Stochastic model of pathogen competition in a descending fitness landscape

To establish the potential of genomic epidemiological models as tools to study evolution, we created a stochastic model using Opqua following and expanding our results from the compartment model (Fig. 3). The model is composed of 500 hosts in a single population, 250 of which start the simulation infected by “wild-type” pathogens with a specific genome sequence of 500 amino acids. Each simulation lasted for 1000 time units, and the result shown for each condition was obtained as the mean of ten replicates. The epidemiological parameters were set to the default values shown for host-host models in Table S1, with the following modifications: a genome length of 500 loci with 20 possible alleles (corresponding to the 20 amino acids abbreviations “ARNDCEQGHILKMFPSTWYV”) at each locus, a mean inoculum of 1 pathogen, a mutation rate of 0.05 1/unit time, and no recombination. The full amino acid alphabet was used in this model to showcase the capabilities of Opqua and genomic epidemiological modeling as a way of describing complex diversity. Since this simulation does not aim to show stochastic tunneling to a specific new fitness peak, all evolutionary trajectories away from wild type are considered of interest, allowing for the use of more complex genomes and allele systems.

The intra-host competitive fitness *φ* of each genome was made to follow the following function, describing an exponential decay in fitness in terms of the relative fitness disadvantage of each successive mutation *λ* (shown on Fig. 3) and the Hamming distance *d* between a mutant genome to the wildtype genome:

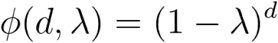

The maximum distance between a mutant genome and wild-type and the mean pairwise distance between genomes at the end of simulation were both computed as Hamming distances.

### Stochastic model of population determinants of evolution

We then used the same modeling framework to examine the effect of population size and migration between populations on evolution across descending fitness landscapes (Fig. 4). The parameters and analyses used were the same as the ones used for the previous stochastic model describing a descending fitness landscape, unless otherwise noted on each figure. The host-host contact rate was fixed to *β*=0.125 1/unit time, and the fitness cost of each successive mutation was set to *λ*=0.5. For the metapopulation model, the system consisted of 10 different interconnected populations of 10 initial hosts each.

### Stochastic model of pathogen determinants of evolution

Finally, to study the effect of different biological factors on evolution across descending fitness landscapes, we modeled a host-vector system using Opqua (Fig. 5). The model consisted of 250 hosts and 250 vectors in a single population, running for 1000 time units in 10 replicates for each condition. Model parameters were set to the default values shown on Table S3, with a few modifications. All simulations had the host-vector contact rate set to 0.125 1/unit time. The fitness cost of each successive mutation was set to *λ*=0.5 within hosts only, and was set to *λ*=0 within vectors, to model the differing selection pressures on the genome in each organism. Simulations which varied recombination rates had a mean number of crossover events in both hosts and vectors set to 5. Other variables were set as shown on Fig. 5, or the default values on Table S2 if not noted.

An analogous set of simulations was repeated in a host-host transmission system as shown in Figs. S5 and S6. All parameters were identical as in the host-vector system described, except for the host-host contact rate set to 0.125 1/unit time and the host-vector contact rate set to zero, as well as the parameters indicated on each figure.

A third set of simulations was carried out using a host-vector transmission model identical to that used for Fig. 5, but this time with separate fitness peaks for genomes in hosts and vectors (Figs. S7, S8). Each fitness peak was devised to be three mutations away from the wild-type sequence, and six mutations from each other peak. The three distinguishing mutations for each peak lie on opposite extremes of the 500-locus wild-type genome, in order to maximize likelihood of recombination. This allows clearer visualization of the effects of recombination in this system. All other parameters were identical as in the host-vector system described for Fig. 5.

## Acknowledgements

The authors would like to gratefully acknowledge Manuela Carrasquilla, Juan Estupiñán, Alejandro Feged Rivadeneira, Felipe González Casabianca, Angélica Knudson, and David Suárez for their valuable insights throughout the development of this project, as well as Christopher V. Turlo and Jelle van der Hilst for feedback on figure design.

## Authors contributions

P.C., V.C. and M.S.V. conceived the idea of the software, designed the study, and edited the manuscript. P.C. implemented the code and wrote the manuscript.

## Data and materials availability

All code necessary to install and use the Opqua modeling framework is available at https://github.com/pablocarderam/opqua. All data and code necessary to reproduce the simulations and analyses of this specific study is available at https://github.com/pablocarderam/fitness_valleys_opqua/.

## Supplementary Materials

**Supplementary Fig. S1.**
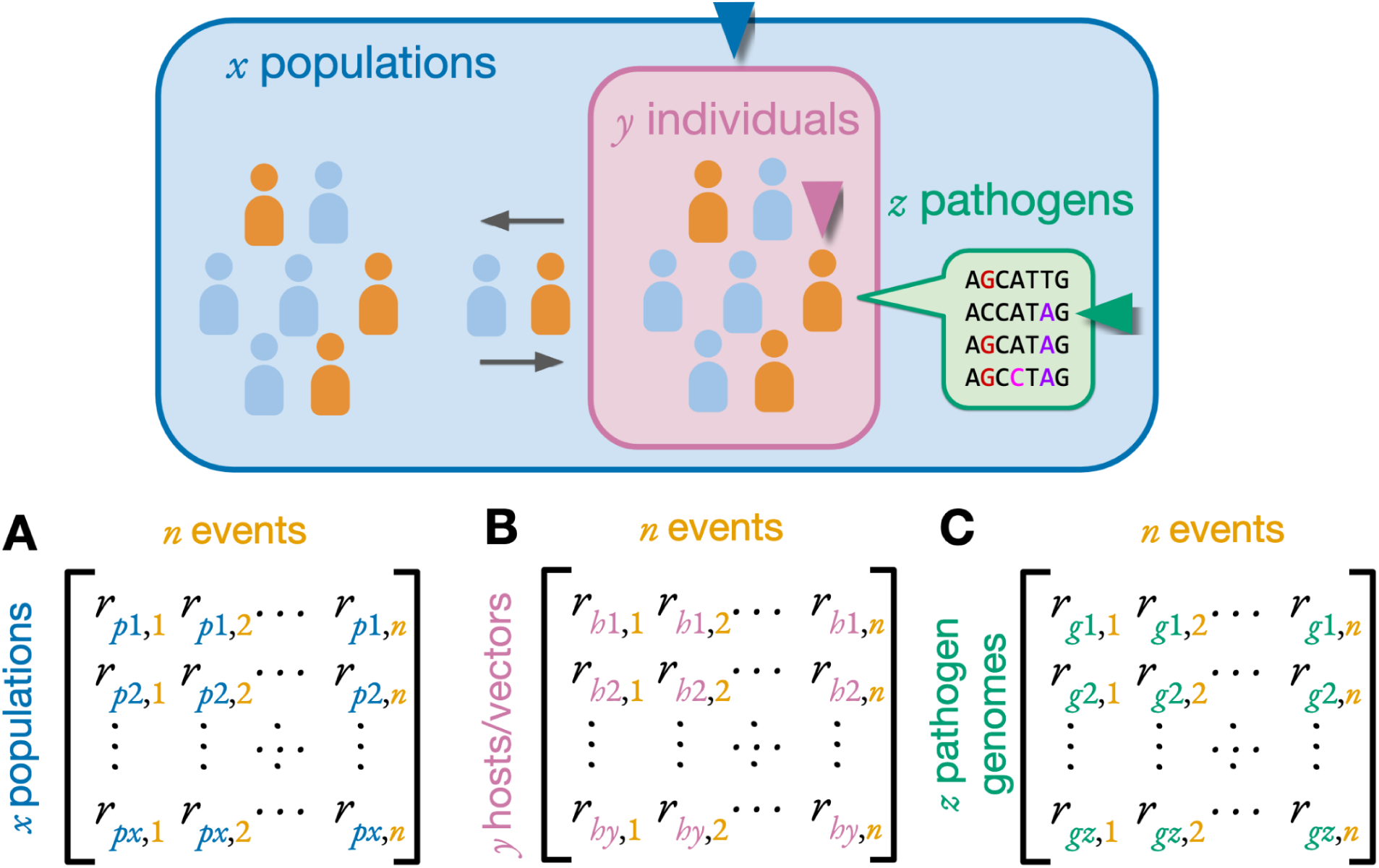
Opqua uses different levels of rate matrices to sample events. **(A)** A matrix contains the rates *r_p_* of *n* different events for *x* different populations within the model. A given event type and population are sampled randomly based on the rate of each event-population combination. **(B)** Each population contains matrices with the event rates *r_i_* for all *y* individual hosts and vectors it contains. A host/vector is sampled randomly based on the rates of the chosen event type for all host/vectors in the chosen population. **(C)** Finally, a similar process occurs within hosts and vectors to randomly sample pathogens within them, based on how their *z* different genomes affect the rates *r_g_* of the chosen event. Some events involve sampling an additional population (migration or inter-population contact), host/vector (inter- or intra-population contact), or pathogen (recombination). Once event type, population(s), host(s) and/or vector(s), and pathogen(s) have been chosen in this manner, the state of the model is adjusted according to the event, and the relevant rate changes are propagated upward from within the pathogens affected to the host(s), vector(s), populations, and overall model they are in.

**Supplementary Fig. S2.**
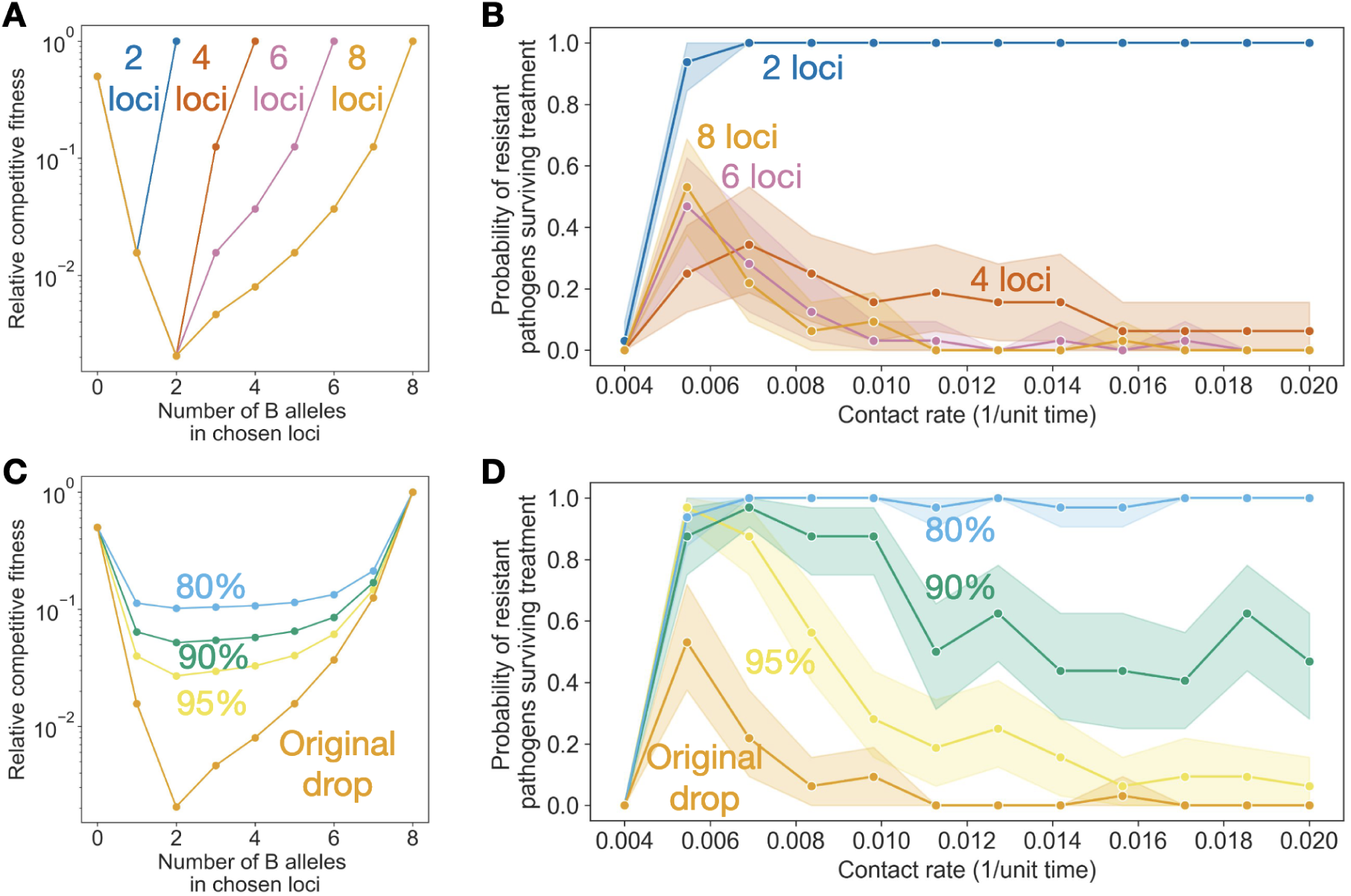
Length and steepness of the fitness valley affect the impact of competition on evolution. We constructed a model to simulate stochastic tunneling of a pathogen across a fitness valley based on genomes with eight bi-allelic loci, as portrayed in Fig. 2. **(A)** By focusing on a subset of loci in each genome, we can vary the length of the fitness valley. **(B)** Longer fitness valleys favor the evolution and survival of pathogens in low transmission environments, as more loci provide more alternative evolutionary paths through the valley but are inhibited by high intra-host competition. In short fitness valleys, the number of paths becomes more restricted while the effect of competition is lessened, favoring environments with increased transmission. **(C)** By adjusting the fitness cost of intermediate mutants, we can vary the steepness and depth of the fitness valley. **(D)** Steep, deep valleys increase the effect of intra-host competition, favoring the evolution and survival of pathogens in low transmission environments more heavily.

**Supplementary Fig. S3.**
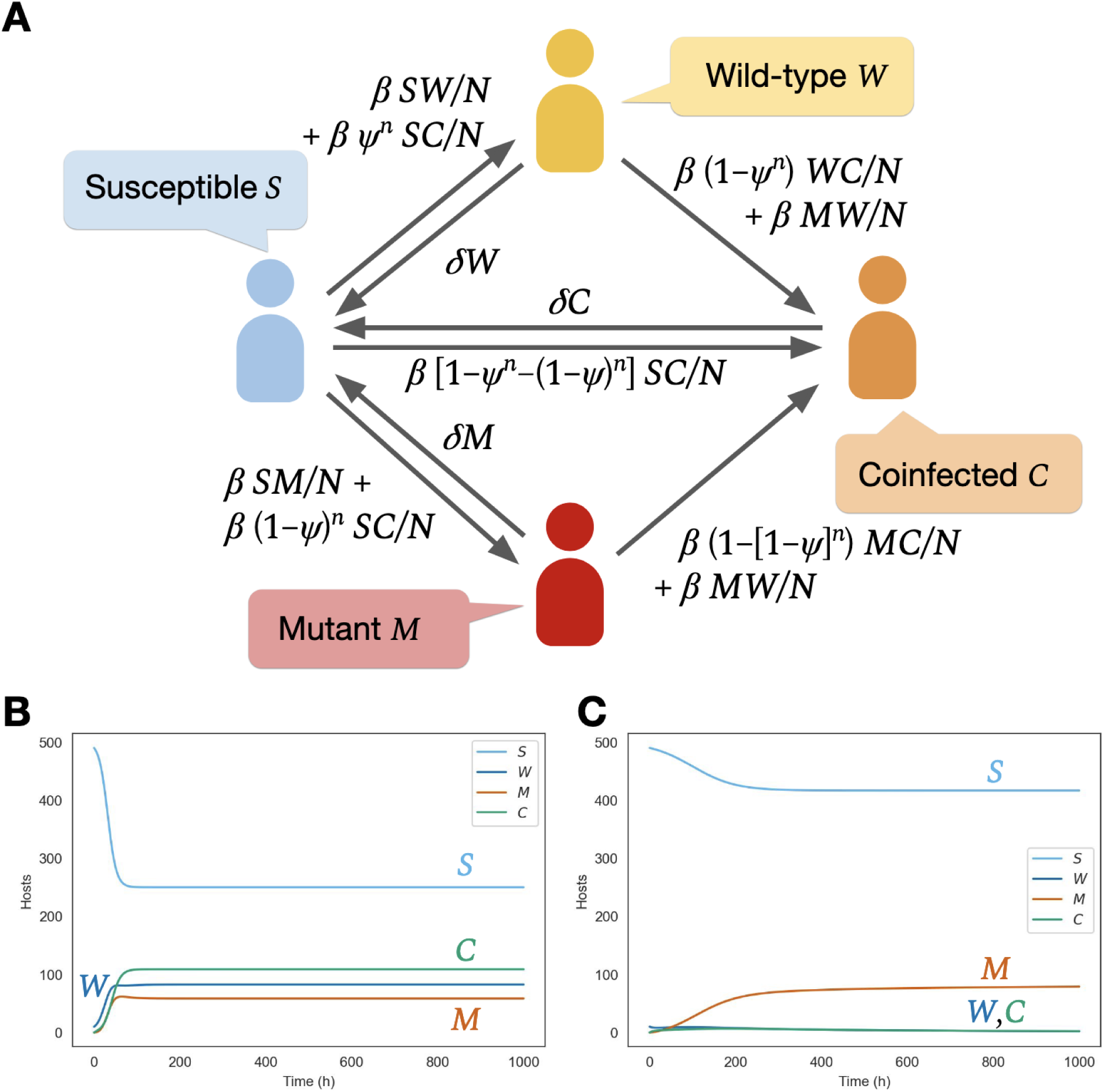
Compartment model structure allows deterministic simulation of dynamics of two strains of pathogens. **(A)** The model consists of compartments for hosts infected with no pathogens (*S*), wild-type pathogens only (*W*), mutant pathogens only (*M*), or coinfected with both kinds of pathogens (*C*). Transitions between compartments are determined by host recovery rate *δ*, contact rate *β*, mutation rates from wild-type to mutants *µ_1_* and vice-versa *µ_2_*, inoculum size *n*, and the probability of wild-type pathogens with higher fitness outcompeting mutants in intra-host competition *ψ*. **(B)** At high transmission intensities (*β=*0.2), unfit mutants (*ψ=*0.67) constitute a small portion of the total population, as shown throughout the rest of this work. **(C)** At low transmission intensities (*β=*0.12), unfit mutants show greater prevalence in the population.

**Supplementary Fig. S4.**
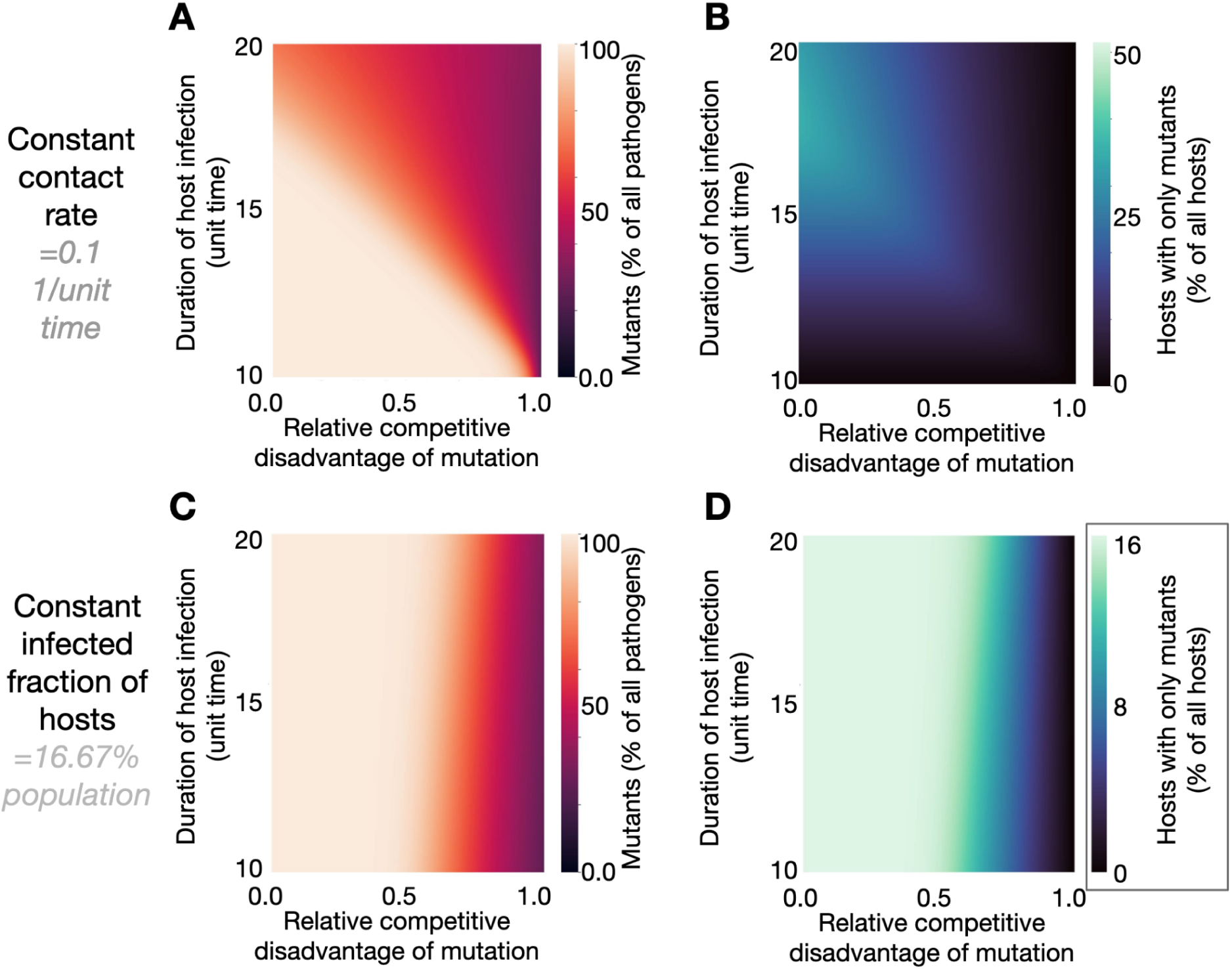
Duration of infection affects mutant fraction primarily through competition for free hosts, as with contact rate. Using the ordinary differential equation model described, we vary the duration of infection (equivalent to 1/recovery rate) while keeping the contact rate constant such that the range of steady-state infected hosts sampled is equivalent to those in Figure 3B and 3C. The resulting **(A)** mutant fractions of pathogens and **(B)** mutant-only fractions of infections are similar to those shown when varying contact rates in Figure 3B and 3C. Small differences in the scale of the effects are due to the fact that at high durations of infection, mutants are removed less frequently through recovery. This can be seen when varying the duration of infection while keeping the fraction of infected hosts constant by simultaneously varying the contact rate. While biologically unlikely to happen, this shows a small increase in fraction of **(C)** mutants among pathogens and **(D)** mutant-only infections among hosts (note change in color scale).

**Supplementary Fig. S5.**
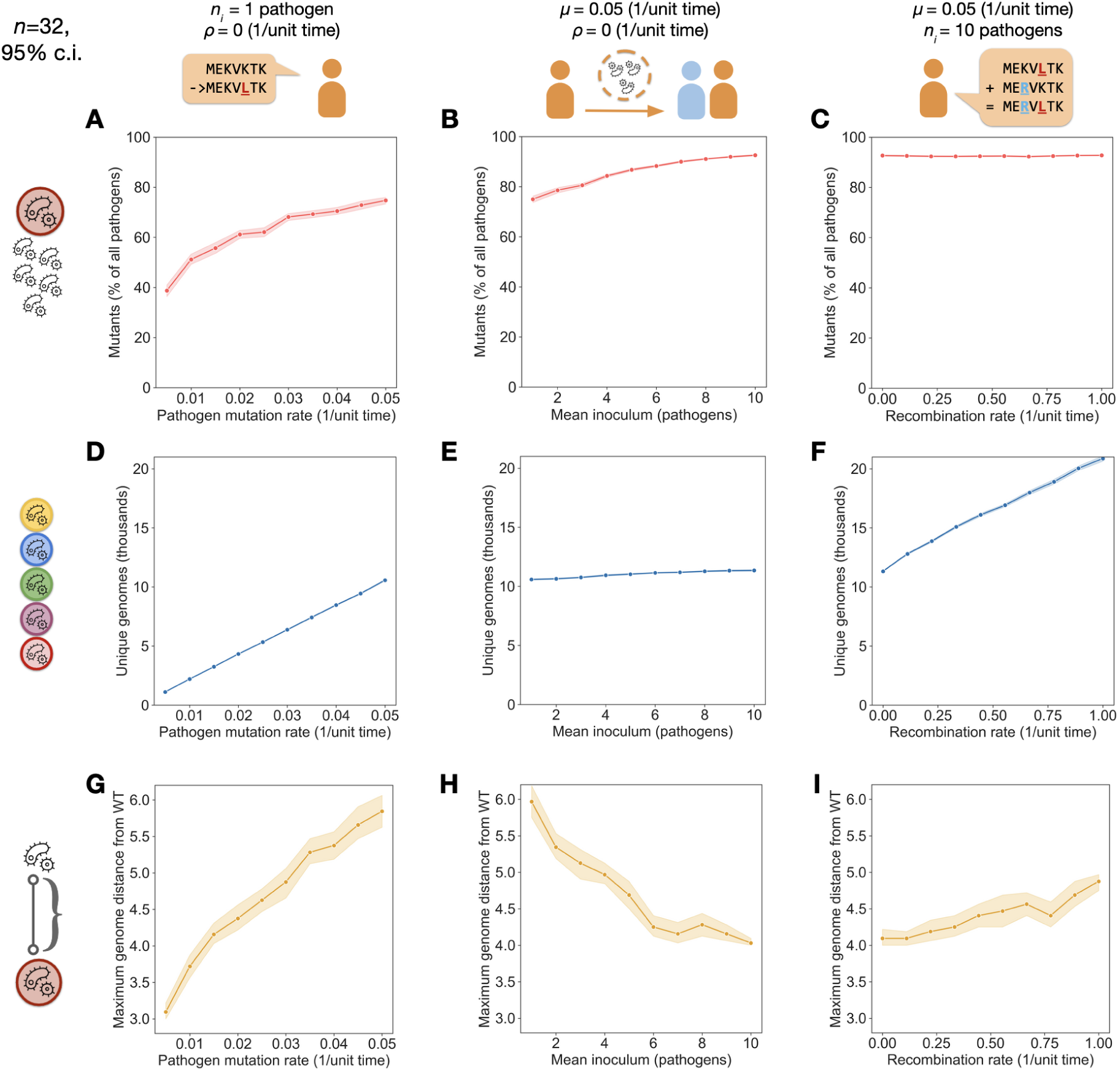
Pathogen biology affects the distribution and dimensions of their evolution for host-host direct transmission. The effects of mutation rate, inoculum size from hosts, and recombination rates are analogous to those observed within hosts for a vector-borne model (Fig. 5). The fraction of mutants in the pathogen population is increased by both **(A)** high mutation rates (*µ*) and **(B)** high mean inoculum sizes (*n_i_*), but is unaffected by **(C)** recombination rates (*ρ*). **(D)** The number of unique pathogen genomes in the simulation, which we treat as a measure of the “width” of evolutionary space explored, increases with high mutation rates, but less so with **(E)** high mean inoculum size. **(F)** Greater recombination increases the number of unique pathogen genomes. Lastly, **(G)** increased mutation rate increases the “depth” of evolutionary space explored by pathogens, measured as the maximum Hamming distance of mutant genomes from the initial wild-type sequence. **(H)** Low inoculum size increases the probability of transmitting mutants without wild-type (WT) competitors, allowing for greater depth in the evolutionary space explored. **(I)** High recombination rates increase depth of evolutionary space, notably without affecting the fraction of mutants in the pathogen population.

**Supplementary Fig. S6.**
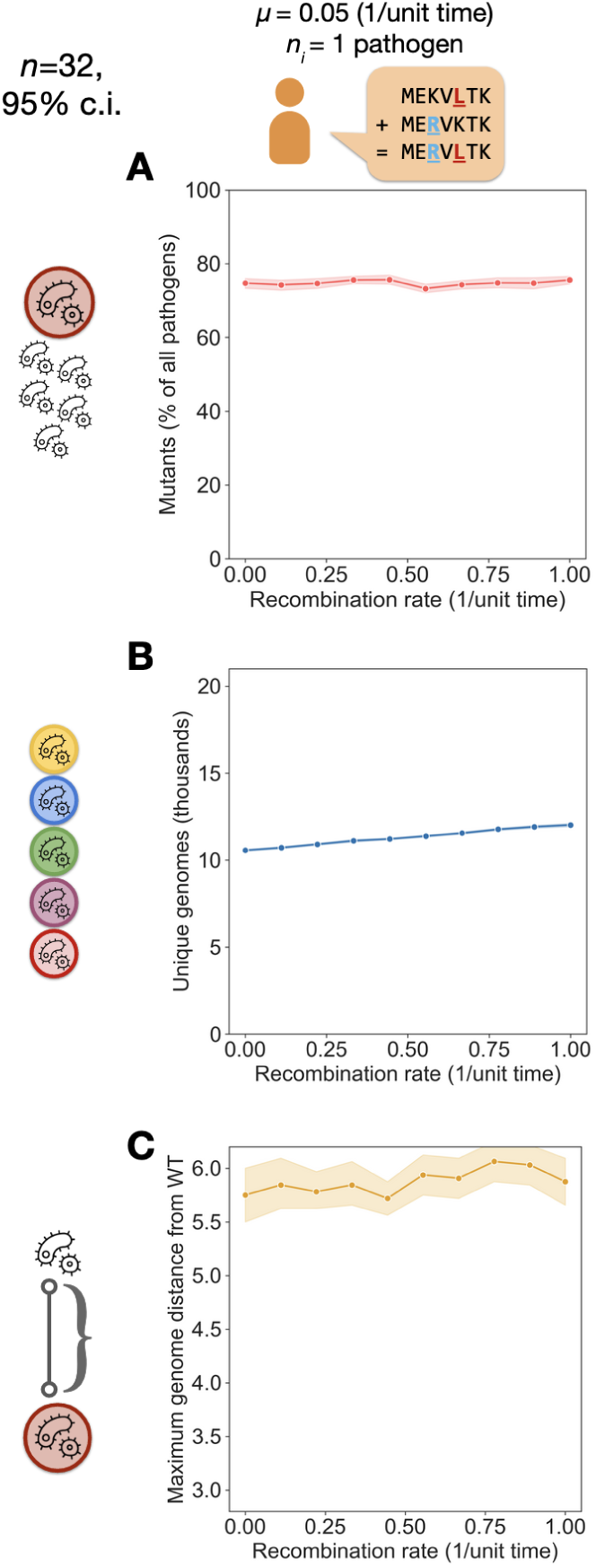
Recombination depends on inoculum size to increase evolutionary distance. By keeping the mutation rate (*µ*) constant and reducing the mean inoculum size (*n_i_*) to 1 in a host-host transmission model with a descending fitness landscape, we can see the effects of recombination on pathogen genome evolution with low inoculums. **(A)** As before, increasing recombination rate does not increase the mutant fraction of pathogens. However, the lower inoculum greatly reduces the effect of recombination on **(B)** the number of unique pathogen genomes explored and **(C)** the maximum Hamming distance from the wild-type pathogen genotype. Neither of these shows increases on the same scale seen for the higher mean inoculum size *n_i_*=10 in Fig. S3.

**Supplementary Fig. S7.**
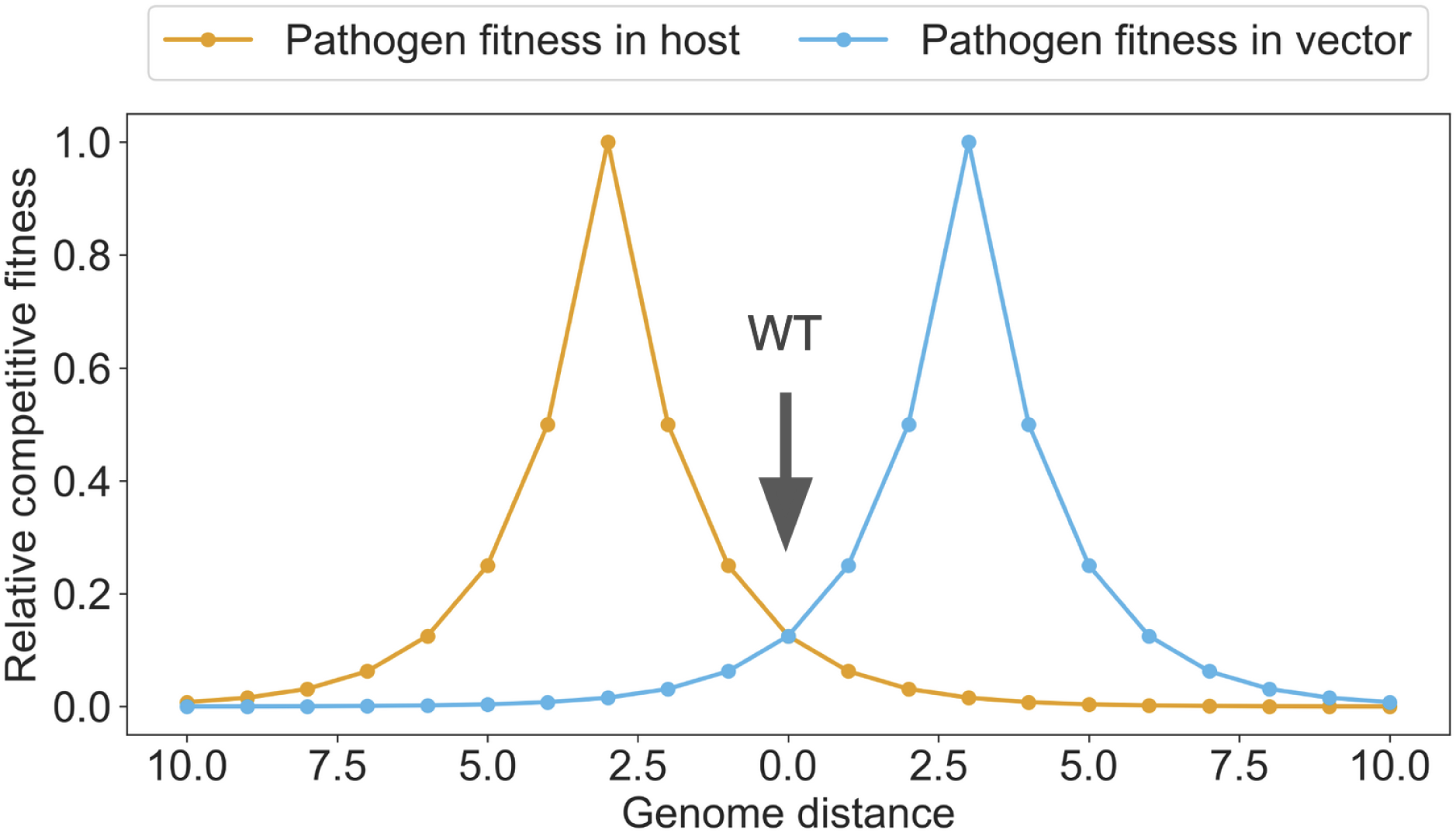
Separate fitness landscapes within hosts and vectors can be used to study pathogen evolution in Opqua. Two fitness functions are devised with exponentially decaying fitness around distinct, optimal genome sequences. These optimal genomes constitute separate fitness peaks for pathogens within hosts and vectors. Each fitness peak is at Levenshtein distance of six mutations from the other. The wild-type (WT) genome sequence used to initiate simulations lies halfway between both peaks, three mutations from each.

**Supplementary Fig. S8.**
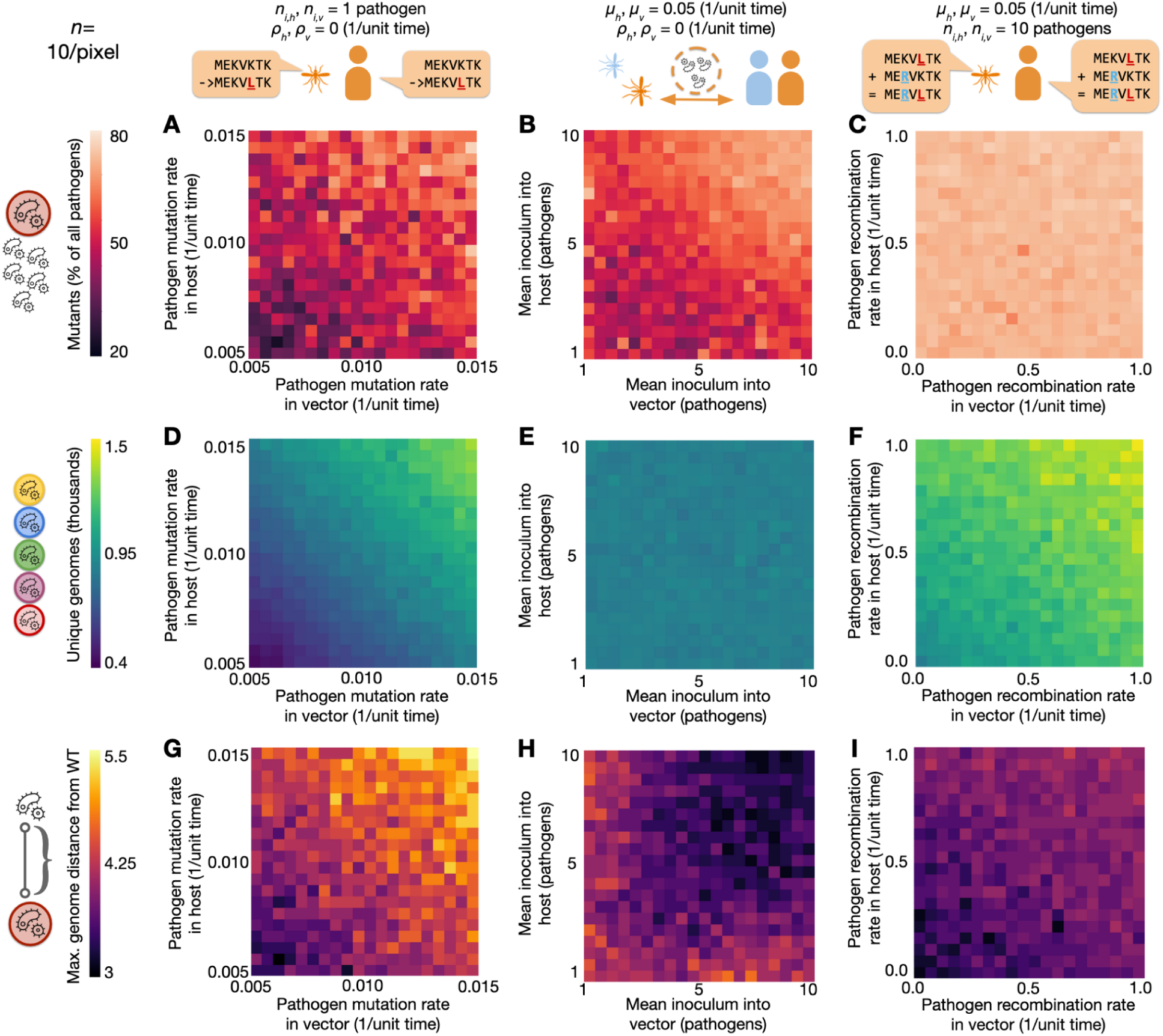
Conflicting fitness landscapes in different life stages generate symmetric distributions of evolution within hosts and vectors. We varied mutation rates, inoculum size, and recombination rates in hosts and vectors as done in the simulations shown on Fig. 5, with the addition of selection within both hosts and vectors (as shown in Fig. S7). The resulting heatmaps show distributions of **(A–C)** mutant pathogens, **(D–F)** unique genomes explored, and **(G–I)** maximum genome distance from the wild-type genome that are symmetric across the diagonal, and correspond to the average of each respective graph on Fig. 5 and its diagonal reflection. This is expected, given the same population bottlenecks described in the main text for the model with only selection in hosts are now present with symmetric selection pressures across the life cycle.

**Table S1:** Default Opqua parameters for host-host transmission models*.

| Parameter name | Default value |
| --- | --- |
| num_loci | 10 |
| possible_alleles | 'ATCG' |
| fitnessHost | (lambda g: 1) |
| contactHost | (lambda g: 1) |
| receiveContactHost | (lambda g: 1) |
| mortalityHost | (lambda g: 1) |
| natalityHost | (lambda g: 1) |
| recoveryHost | (lambda g: 1) |
| migrationHost | (lambda g: 1) |
| populationContactHost | (lambda g: 1) |
| receivePopulationContactHost | (lambda g: 1) |
| mutationHost | (lambda g: 1) |
| recombinationHost | (lambda g: 1) |
| fitnessVector | (lambda g: 1) |
| contactVector | (lambda g: 1) |
| receiveContactVector | (lambda g: 1) |
| mortalityVector | (lambda g: 1) |
| natalityVector | (lambda g: 1) |
| recoveryVector | (lambda g: 1) |
| migrationVector | (lambda g: 1) |
| populationContactVector | (lambda g: 1) |
| receivePopulationContactVector | (lambda g: 1) |
| mutationVector | (lambda g: 1) |
| recombinationVector | (lambda g: 1) |
| contact_rate_host_vector | 0 |
| transmission_efficiency_host_vector | 0 |
| transmission_efficiency_vector_host | 0 |
| contact_rate_host_host | 2.00E-01 |
| transmission_efficiency_host_host | 1 |
| mean_inoculum_host | 1.00E+01 |
| mean_inoculum_vector | 0 |
| recovery_rate_host | 1.00E-01 |
| recovery_rate_vector | 0 |
| mortality_rate_host | 0 |
| mortality_rate_vector | 0 |
| recombine_in_host | 1.00E-04 |
| recombine_in_vector | 0 |
| num_crossover_host | 1 |
| num_crossover_vector | 0 |
| mutate_in_host | 1.00E-06 |
| mutate_in_vector | 0 |
| death_rate_host | 0 |
| death_rate_vector | 0 |
| birth_rate_host | 0 |
| birth_rate_vector | 0 |
| vertical_transmission_host | 0 |
| vertical_transmission_vector | 0 |
| inherit_protection_host | 0 |
| inherit_protection_vector | 0 |
| protection_upon_recovery_host | None |
| protection_upon_recovery_vector | None |

\*For a detailed description of each parameter, consult the documentation at https://github.com/pablocarderam/opqua#newsetup

**Table S2:**
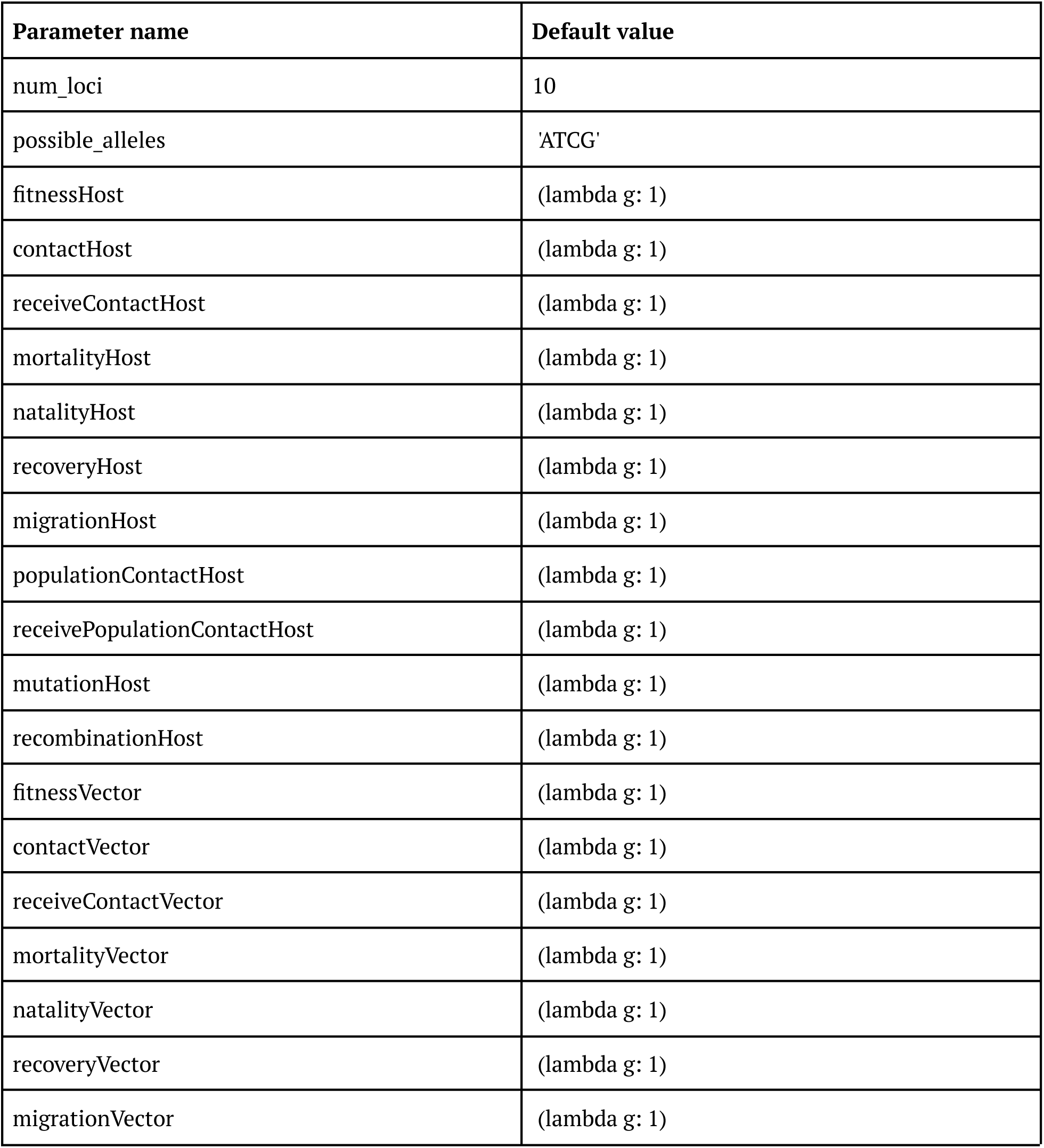
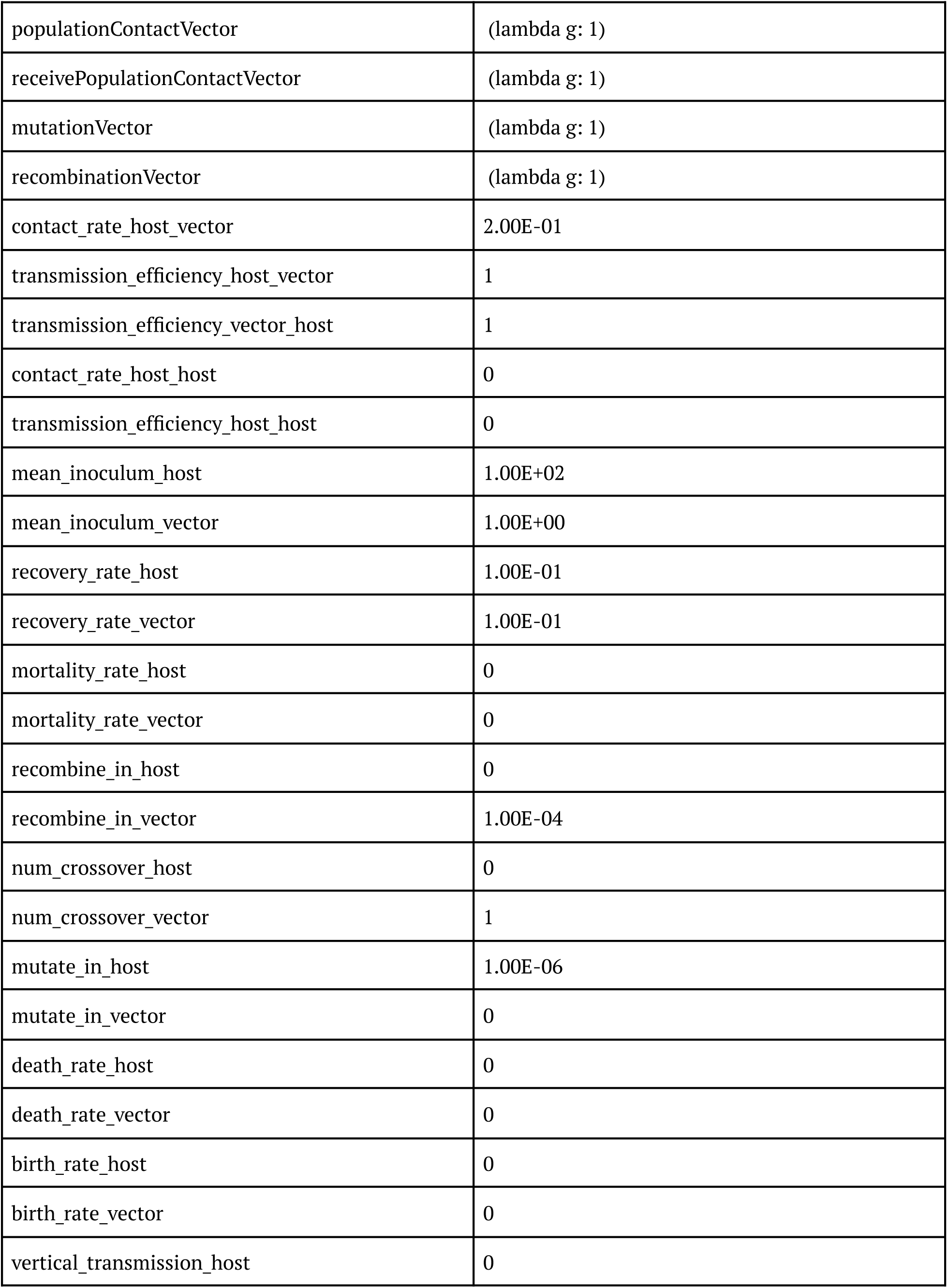
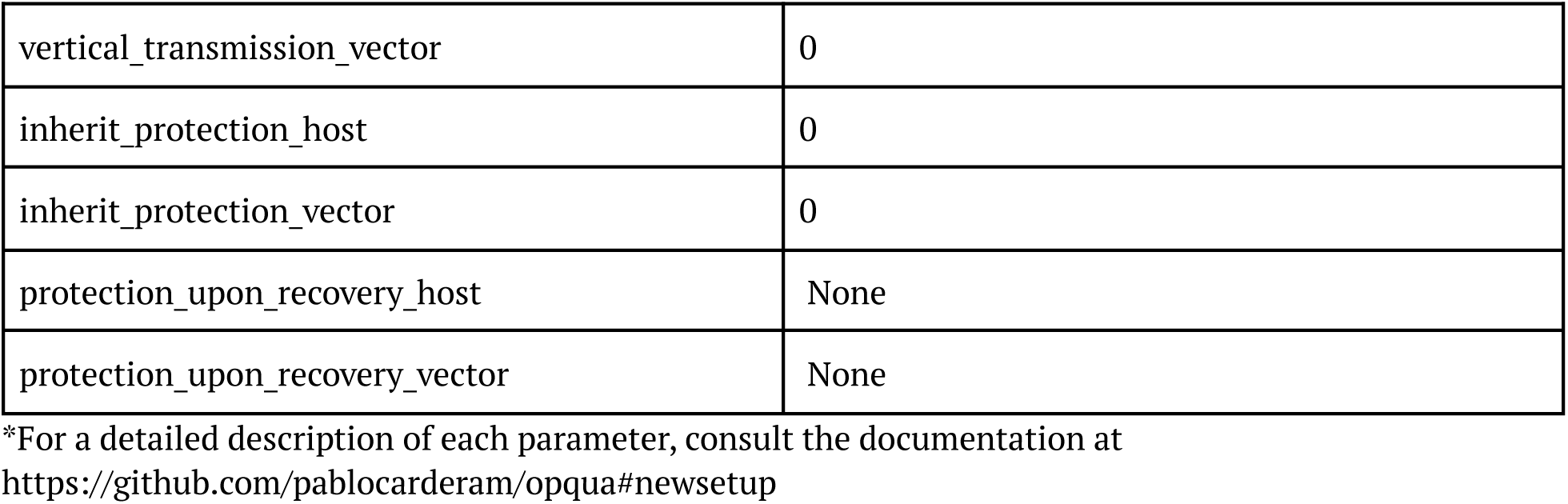
Default Opqua parameters for vector-borne transmission models*.

**Table S3:** Parameters used for two-strain compartment model.

| Parameter name | Description | Value |
| --- | --- | --- |
| $\delta$ | host recovery rate | 1.00E-01* |
| $\beta$ | host contact rate | 1.10E-01* |
| $\varphi$ | probability of wild-type pathogens outcompeting mutants in intra-host competition | 1/(2-0.1)* |
| $n_i$ | size of inoculum | 1 |
| $\mu_1$ | mutation rate from wild-type to mutant | 5.00E-02 |
| $\mu_2$ | mutation rate from mutant to wild-type | 0 |
| $N$ | total population size | 5.00E+02 |

\*Parameters vary according to simulation

